# Multiplexed cytokine and antigen mRNA administration generates durable anti-tumor immunity against pancreatic cancer

**DOI:** 10.1101/2025.10.13.681939

**Authors:** Chaitanya N. Parikh, Kelly D. DeMarco, Griffin I. Kane, Nikita Bhalerao, Ronnie W. Dinnell, Boyang Ma, Hadiya K. Giwa, Zhen Zhao, Lin Zhou, Katherine C. Murphy, Loretah Chibaya, Haruka Mori, Youwei Qiao, Brian C. Lewis, Wen Xue, Jason R. Pitarresi, Prabhani U. Atukorale, Marcus Ruscetti

## Abstract

Pancreatic ductal adenocarcinoma (PDAC) remains a devastating malignancy characterized by limited therapeutic options for advanced disease. Immunotherapy, in particular, has had dismal success rates in the PDAC due to a tumor microenvironment (TME) that contributes to immune exclusion and poor drug delivery. Many cytokines necessary for Natural Killer (NK) and T cell chemotaxis, activation, and cytotoxicity are absent in the PDAC TME. Despite their early success, cytokine therapies have largely failed in the treatment of solid tumors as a result of the lack of efficacy of single cytokine administration and toxicities from systemic delivery. To overcome these limitations, we designed multiplexed mRNA cocktails encoding diverse interleukins, chemokines, and interferons for intratumoral delivery. Administration of a cytokine-encoding mRNA mixture into mice with orthotopically transplanted PDAC tumors achieved robust yet transient cytokine expression locally in the PDAC TME, leading to NK cell and CD8^+^ T cell immunity and reduced tumor growth and fibrosis in multiple mouse models. Combining cytokine mRNAs with those encoding tumor-associated antigens further activated CD8^+^ T cell-mediated tumor control and enhanced survival after just a single dose in PDAC-bearing mice. Remarkably, lipid-based nanoparticle (NP) encapsulation of an all-in-one cytokine and antigen mRNA cocktail allowed safe systemic administration and local delivery of these immunogenic signals to autochthonous PDAC tumors in genetically engineered mouse models, culminating in complete tumor responses in 50% of animals. These results suggest that multiplexed mRNA approaches to delivering cytokine signals and antigens generally absent in the TME could pave the way for an effective immunotherapy for PDAC.

## INTRODUCTION

Pancreatic ductal adenocarcinoma (PDAC) is a highly aggressive malignancy with a median 5-year survival rate of only 13%, and is projected to be the second leading cause of cancer-related deaths in the United States by 2030^1,2^. Most patients present with disseminated disease and are not eligible for surgical resection^3^. Standard-of-care chemotherapy regimens have limited efficacy, in part due to a fibrotic and hypovascular tumor microenvironment (TME) that leads to poor drug delivery and uptake^4–7^. Moreover, rampant immune suppression and exclusion of cytotoxic Natural Killer (NK) and T lymphocytes in the PDAC TME, coupled with weak tumor immunogenicity, contribute to *de novo* resistance to immune checkpoint blockade (ICB) inhibitors targeting PD-1/PD-L1 and/or CTLA-4 that have demonstrated remarkable success in treating other chemo-resistant malignancies such as non-small cell lung cancer (NSCLC) and melanoma^8–10^. Hence, there is a pressing need for innovative therapeutic strategies to tackle the suppressive TME of PDAC and achieve durable tumor control.

*KRAS* mutations are present in >90% of PDAC lesions and drive disease onset and progression intrinsically by promoting tumor cell growth and survival and extrinsically through remodeling the surrounding TME to mediate immune evasion^11^. We have previously shown that targeting signaling downstream of KRAS with a combination of the MEK inhibitor trametinib and CDK4/6 inhibitor palbociclib (T/P) can promote tumor vascularization and CD8^+^ T cell accumulation in the TME, leading to increased uptake and sensitivity to chemotherapy and anti-PD-1 ICB^12^. These anti-tumor effects were mediated in part through the induction of cellular senescence and its accompanying senescence-associated secretory phenotype (SASP), a collection of hundreds of pleiotropic cytokines, growth and angiogenic factors, matrix metalloproteinases, and lipid species that can remodel the TME and in particular immune responses in dynamic ways^13^. Indeed, combining these senescence-inducing therapies with EZH2 inhibitors or lipid nanoparticle (NP)-loaded STING and TLR4 agonists further enhanced the pro-inflammatory cytokine arm of the SASP, producing robust anti-tumor NK and CD8^+^ T cell immunity that led to even complete responses in preclinical PDAC mouse models^14,15^. Blockade of different SASP factors revealed key interleukins (IL-12,-15,-18), interferons (IFNβ), and chemokines (CCL2, CXCL9/10/11) necessary for anti-tumor NK and T cell immunity in PDAC following senescence induction. These findings suggest that the direct delivery of SASP-related cytokines to tumors may be a novel strategy to reactivate immunity against PDAC.

Cytokines are a diverse group of small proteins that act as signaling molecules to mediate the recruitment, proliferation, and activation of various immune cells^16^. As cytokine expression is frequently altered in human cancers, cytokine administration or blockade has long been pursued as a therapeutic approach to immunologically treat tumors^17^. Dating back to the 1980/90’s, systemic delivery of recombinant cytokine proteins IFNα or IL-2 that can stimulate innate and adaptive immune responses were some of the first approved immunotherapies for treating immunologically “hot” cancers like melanoma and renal cell carcinoma^18–21^. However, the modest efficacy of single cytokine administration combined with the often severe toxicities elicited by high-dose systemic delivery have limited the clinical utility of these and other cytokines such as IFN*γ* and IL-12^22–26^.

Advances in mRNA therapeutics and drug delivery methods in recent years have offered the opportunity to revisit cytokine therapy for cancer. Indeed, work from a number of groups has shown that multiple cytokine-encoding mRNAs can be delivered intratumorally to elicit potent anti-tumor immune responses without systemic toxicities in melanoma and other immunologically “hot” tumor models, even outperforming recombinant protein approaches^27–29^. A recent study demonstrated that IL-12 encoding mRNAs could be administered intratumorally into PDAC-bearing mice to reprogram immune suppressive myeloid cells and activate T cell immunity^30^. This suggests that mRNA-based cytokine strategies could also be applied to remodel the TME of immune suppressed cancers such as PDAC, where drug delivery is a major limitation. Here, we established an *in vitro* transcription pipeline for *de novo* mRNA synthesis and multiplexing to produce multiple interleukins, IFNs, and chemokines that we have previously demonstrated to be important for mediating both innate and adaptive anti-tumor immunity in preclinical PDAC models^12,14,15^. Using preclinical transplant and autochthonous mouse models of pancreatic cancer, we tested whether simultaneous administration of different cytokine combinations as free mRNAs intratumorally or via systemic delivery through NP encapsulation could reactivate immune surveillance and achieve durable tumor control.

## RESULTS

### Intratumoral delivery of cytokine-encoding mRNA cocktail drives safe and robust local cytokine expression in orthotopic PDAC models

We established an *in vitro* transcription (IVT) pipeline to synthesize mRNAs in multiplex encoding diverse cytokines we have previously demonstrated to be necessary for anti-tumor NK and T cell immunity following therapy-induced senescence in preclinical PDAC models^12,14,15^, including the interleukins IL-12, IL-15, and IL-18, chemokines CCL5 and CXCL10, and the Type I interferon IFNβ (Fig. 1a). mRNAs were modified with N1-methylpseudouridine triphosphate (m¹ΨTP) during IVT to enhance translational efficiency in mammalian systems and minimize activation of non-specific innate immune responses, and underwent a cellulose-based purification strategy to remove dsRNA contaminants that can produce off-target effects as previously described^31–33^. We confirmed the successful synthesis of cellulose-purified mRNA on agarose gel (Extended Data Fig. 1a). To validate their biological activity and confirm appropriate protein production and secretion, purified mRNAs were transfected into PDAC tumor cell lines derived from PDAC tumors in *Pdx1-Cre; Kras^LSL-G12D/wt^;Trp53^LSL-R172H/wt^* (*KPC*) genetically engineered mouse models (GEMMs), as well as cancer-associated fibroblast (CAF) lines derived from PDAC tumors in *Pdx1-Cre; Kras^LSL-G12D/wt^*; Trp53*^LSL-R172H/wt^*; Rosa26^LSL-YFP/LSL-YFP^ *(KPCY)* GEMMs^34,35^. Lipofectamine transfection of 6 cytokine (IL-12/-15/-18, CCL5, CXCL10, IFN-β) mRNAs simultaneously *in vitro* yielded a significant and robust increase in secreted amounts of all cytokines 24 hours post-transfection in tumor cells and to an even greater degree in CAFs that are highly prevalent in the desmoplastic PDAC TME (Extended Data Fig. 1b,c). The level of secretion of many of these cytokines after mRNA transfection was an order of magnitude higher compared to their quantities following treatment with trametinib and palbociclib (T/P), which we have previously demonstrated could induce their expression as part of an NF-*κ*B-driven SASP (Extended data Fig. 1d). This highlights the potential of this mRNA approach to produce a pro-inflammatory cytokine milieu in the PDAC TME.

**Fig 1.**
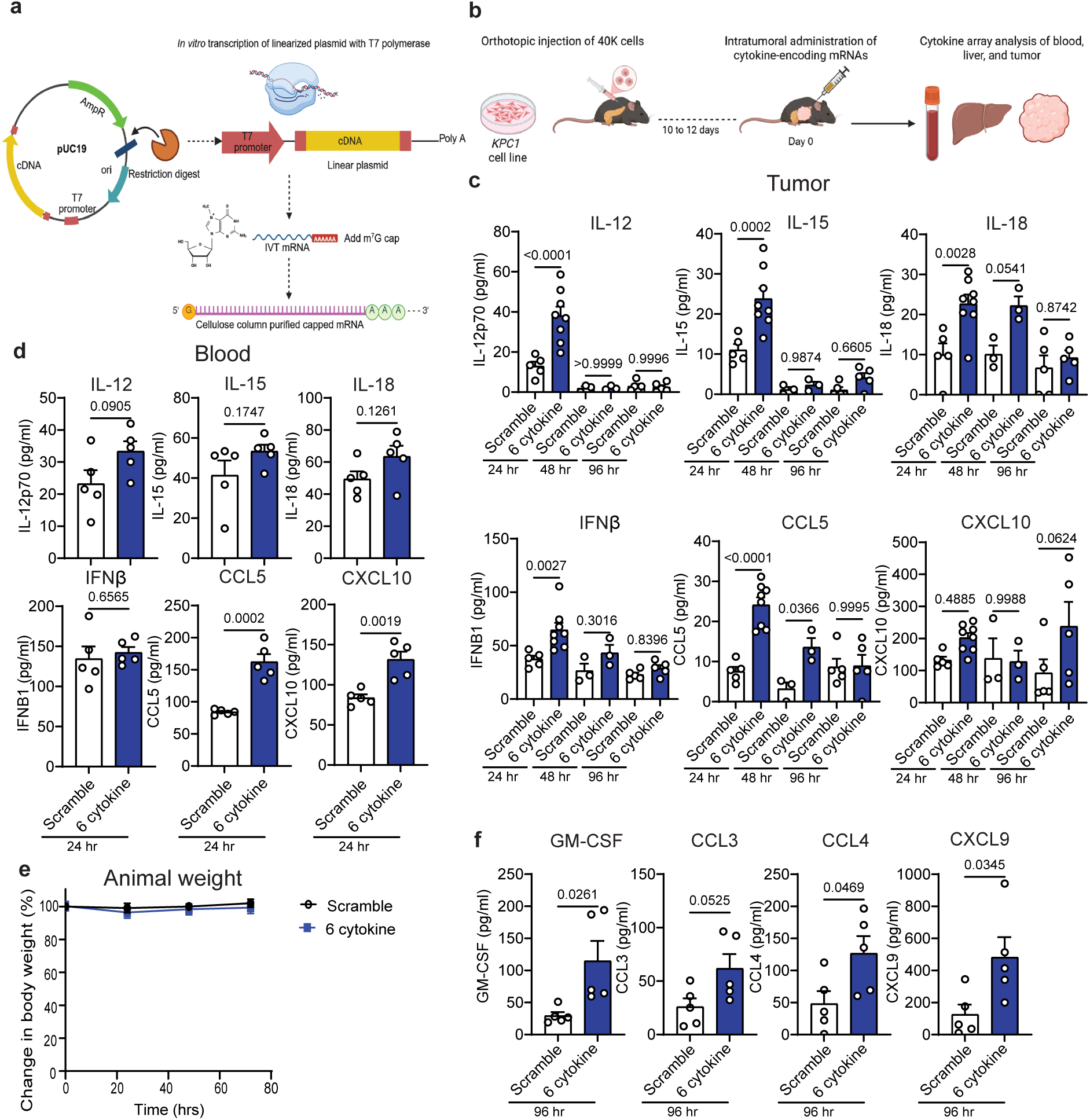
Cytokine mRNA administration drives safe and robust local cytokine expression following intratumoral injection into orthotopic PDAC models. **a**, Schematic of mRNA synthesis pipeline. **b**, *KPC1* PDAC tumor cells expressing luciferase-GFP were orthotopically injected into the pancreas of 8-to 12-week-old female C57BL/6 mice. Following tumor establishment, mice received intratumoral injections of 60 µg of either scramble mRNA or a 6-cytokine mRNA cocktail (10 µg each of IL-12, IL-15, IL-18, IFNβ1, CCL5, and CXCL10). Tumor, liver, and blood samples were collected at 24-, 48-, and 96-hours (hrs) post-injection for cytokine array analysis. **c,d**, Cytokine array results from tumor tissues (c) and blood (d) collected at indicated time points following intratumoral mRNA administration as described in b (*n* = 3–8 mice per group). **e**, Change in weight of *KPC1* tumor-bearing mice at 24-, 48-, and 72 hrs following one intratumoral dose of either scramble mRNA (60 µg) or a 6-cytokine mRNA cocktail (10 µg each of IL-12, IL-15, IL-18, IFNβ1, CCL5, and CXCL10; 60 µg total) (*n* = 5 mice per group). f, Cytokine array results from tumor tissues collected at 96 hrs post-intratumoral mRNA administration as described in b (*n* = 5 mice per group). *P* values in c were calculated using ordinary one-way ANOVA with Sidák’s correction, and those in d and f using two-tailed, unpaired Student’s t-test. Error bars are mean ± s.e.m.

We next evaluated the capacity of these mRNAs to produce secreted cytokines within PDAC tumors in C57BL/6 mice orthotopically transplanted with *KPC1* tumor cells. Once tumors in the pancreas were confirmed by ultrasound imaging (10-12 days post-surgery), the cocktail of 6 free (naked) cytokine-encoding mRNAs (*Il12/15/18, Ccl5, Cxcl10, Ifnb1*) at 10 µg per mRNA, or 60 µg of scrambled mRNA as a control, was administered intratumorally (Fig. 1b). Cytokine array analysis demonstrated increased quantities of each of the cytokines encoded by their corresponding mRNAs within the tumor milieu 24 hours post-injection (Fig. 1c). 10 µg of free *Il12* mRNA produced a similar induction of IL-12 protein levels as either 60 µg of free mRNA or 10 µg of lipid nanoparticle (NP) encapsulated mRNA delivered intratumorally (Extended Data Fig. 1e,f), demonstrating that the low doses of free mRNA we are using are sufficient for cytokine protein production *in vivo*. This pulse in cytokine levels was transient, as their expression was reduced after 48 hours and returned to baseline for most cytokines 96 hours after mRNA dosing (Fig. 1c). With the exception of chemokines CCL5 and CXCL10 that act through gradients to promote immune chemotaxis, levels of other cytokines did not change significantly in the blood or systemically in the liver after intratumoral cytokine mRNA administration (Fig. 1d and Extended Data Fig. 1g). Importantly, treatment was well-tolerated, with no significant changes in mouse body weight observed in mice treated with cytokine mRNAs as compared to control Scramble mRNAs (Fig. 1e).

Broader cytokine array analysis at 96 hours post-injection revealed altered levels of other cytokines not engineered in the administered mRNA cocktail (Extended Data Fig. 1h). In particular, we observed a significant increase in expression of GM-CSF, a key cytokine involved in dendritic cell (DC) differentiation and pro-inflammatory responses^36,37^, as well as chemokines CCL3, CCL4, and CXCL9 that are important drivers of NK and T cell recruitment and associated with better anti-PD-1 immunotherapy responses^38–40^ (Fig. 1f). On the other hand, other pro-inflammatory cytokines known to be induced by endogenous mRNAs through activating Toll-like receptors (IL-1α, IL-1β, IL-6, TNFα) were not significantly altered by injection of our cytokine mRNAs engineered with synthetic nucleoside modifications^33,41^ (Extended Data Fig. 1h). Together, these findings demonstrate that intratumoral administration of a low dose cytokine-encoding mRNA cocktail not only safely augments local expression of the delivered cytokines, but also induces a secondary wave of cytokines that could potentially amplify anti-tumor immune responses.

### mRNA multiplexing uncovers optimal cytokines to enhance tumor immunogenicity and activate innate and adaptive immune responses in PDAC

Given the modularity of our mRNA synthesis pipeline, we set out to investigate the contribution of individual cytokines to different arms of the immune response to determine the optimal combination to mediate effective anti-tumor immunity in PDAC. One the first hurdles to anti-tumor immunity is the inability of cytotoxic lymphocytes such as NK and CD8^+^ T cells to infiltrate into the PDAC TME. As such, we first performed *in vitro* transwell migration assays with NK or CD8^+^ T cells purified from mouse spleens in the top chamber and conditioned media from *KPC* PDAC tumor cells transfected with different single or combinatorial mRNA formulations (or control Scramble mRNAs) in the bottom chamber to assess the impact of cytokines on lymphocyte chemotaxis (Fig. 2a). While individual cytokine mRNAs tested (IL-12, IL-18, CCL5, CXCL10) were not sufficient on their own to promote NK cell chemotaxis, the combination of all six significantly increased NK cell migration *in vitro* compared to control Scramble mRNA (Fig. 2b). IL-18 and CCL5 were necessary for NK cell chemotaxis, as their omission from the 6 cytokine mRNA cocktail reversed its NK cell migratory effects. On the other hand, IL-15 was dispensable, with the resulting 5 cytokine combination sufficient to promote NK cell migration (Fig. 2b). Similarly, a combination of 4 mRNAs encoding cytokines IL-12, IL-18, IFN-β, and CXCL10 produced significantly increased CD8^+^ T cell migration through a transwell insert, with again IL-15 and now CCL5 being unnecessary (Fig. 2c).

**Fig 2.**
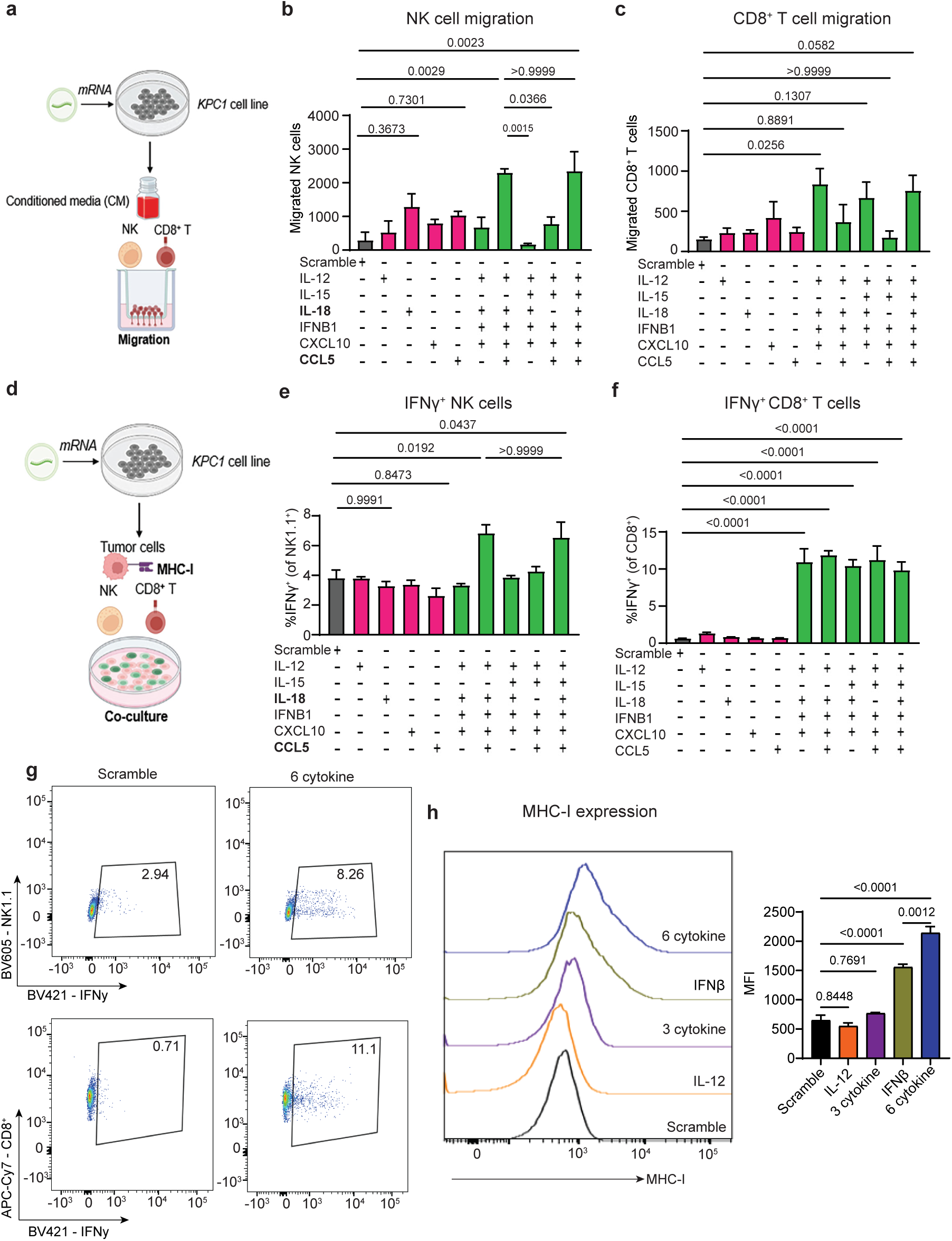
Multiple classes of cytokine mRNAs are necessary to optimally induce NK and CD8^+^ T cell migration and activation and enhance tumor antigen presentation *in vitro*. **a**, Schematic of *ex vivo* NK and CD8⁺ T cell migration assays using conditioned media from *KPC1* PDAC tumor cells 24 hrs after transfection with scramble (3 µg) or cytokine-encoding mRNAs (0.5 µg of each) alone or in combination. **b**,**c**, Quantification of number of NK cells **(b)** and CD8⁺ T cells **(c)** that migrated toward conditioned media from *KPC1* cells transfected with different mRNAs as in **a** (*n* = 3 independent samples per group). **d**, Schematic of NK and CD8⁺ T cell co-culture assays with *KPC1* PDAC tumor cells transfected with mRNAs as in **a**. **e**,**f**, Flow cytometry analysis of IFNγ positivity in NK cells **(e)** and CD8⁺ T cells **(f)** following co-culture with *KPC1* cells transfected with different mRNAs as in **d** (*n* = 3 independent samples per group). **g**, Representative flow cytometry plots showing IFNγ expression in CD45^+^ CD3^-^NK1.1^+^ NK cells (top) and CD45^+^ CD3^+^ CD8^+^ T cells (bottom) following co-culture with *KPC1* cells transfected with indicated mRNAs as in **d. h**, Representative histograms (left) and quantification of mean fluorescent intensity (MFI) of MHC-I (H-2kb) expression on *KPC1* cells following transfection with either scramble mRNA (3 µg), IL-12 or IFNβ1 mRNA alone (0.5 µg), a cocktail of 3-cytokines including IL-12, CCL5, and CXCL10 (0.5 µg each; 1.5 µg total), or a cocktail of 6-cytokines including IL-12, IL-15, IL-18, IFNβ1, CCL5, and CXCL10 (0.5 µg each; 3 µg total). *P* values in **b**,**e**, and **h** were calculated using ordinary one-way ANOVA with Tukey’s correction, and those in **c**,**f** were calculated using ordinary one-way ANOVA with Dunnett’s correction. Error bars are mean ± s.e.m.

We next co-cultured murine NK or CD8^+^ T cells with mRNA-transfected *KPC* PDAC tumor cells and assessed the impact on their effector functions as measured by IFNγ production (Fig. 2d). Flow cytometry analysis of NK cells revealed the combination of all 6 cytokines, or 5 cytokines without IL-15, was required for enhanced IFNγ positivity (Fig. 2e,g), matching the cytokine requirements we uncovered for NK cell migration *in vitro*. For CD8^+^ T cells, the combination of 4 mRNAs encoding cytokines IL-12, IL-18, IFN-β, and CXCL10 was sufficient to produce significantly increased CD8^+^ T cell IFNγ production (Fig. 2f,g). Collectively, these results demonstrate that multiple mRNA cytokines are needed to enhance NK and T cell chemotaxis and effector functions, and that IL-15 is dispensable for these phenotypes.

Antigen-dependent CD8^+^ effector T cell responses rely on the expression and recognition of antigens expressed on the surface of tumor cells through major histocompatibility complex Class I (MHC-I) molecules, which are generally lowly expressed in PDAC^42,43^. To investigate if cytokine mRNAs could also enhance pancreatic tumor cell immunogenicity through upregulating MHC-I, *KPC* cells were transfected with individual mRNAs encoding IL-12 and IFN-β that have established roles^44,45^ in inducing MHC-I alone or as part of the full six cytokine mRNA cocktail. MHC-I surface expression was assessed by flow cytometry 24 hours post-transfection. Whereas IL-12 mRNAs had no impact on MHC-I levels, IFN-β mRNA significantly enhanced MHC-I expression on tumor cells (Fig. 2h). The full 6 cytokine mRNA combination further increased MHC-I intensity compared to IFN-β mRNA treatment alone, demonstrating that use of the cytokine combinations can further enhance antigen presentation. Still, mRNA combinations lacking IFN-β (3 cytokine with IL-12, CXCL10, CCL5) failed to induce MHC-I upregulation, underscoring its essential role (Fig. 2h). Collectively, these findings provided strong rationale for the delivery of a combination of mRNAs encoding ILs, IFNs, and chemokines to fully elicit NK and CD8^+^ T cell migration and activation and antigen presentation required to achieve anti-tumor immunity against PDAC.

### Intratumoral administration of cytokine mRNA cocktail mobilizes innate and adaptive immunity and produces anti-tumor and fibrotic effects in PDAC-bearing mice

We next aimed to determine if the cytokine mRNA cocktail could alter immune responses in the PDAC TME *in vivo* in mice bearing orthotopically transplanted *KPC1* tumors that we have previously shown to be immunologically “cold”^12,14,15^. Given the transient nature of cytokine expression following intratumoral mRNA administration (Fig. 1c), we conducted repeated intratumoral injections of our cytokine mRNA cocktail every three days to enable long-term assessment of the immune response over a 12 day period (Fig. 3a). In addition to some mice receiving 10 μg each of IL-12, IL-15, IL-18, IFN-β, CXCL10, and CCL5 mRNAs (6 cytokine cocktail), a separate cohort were also injected intratumorally with 10 μg of IL-12 mRNA alone for comparison, given recent reports demonstrating that IL-12-encoding mRNAs can activate immune responses in PDAC mouse models^30,46^.

**Fig 3.**
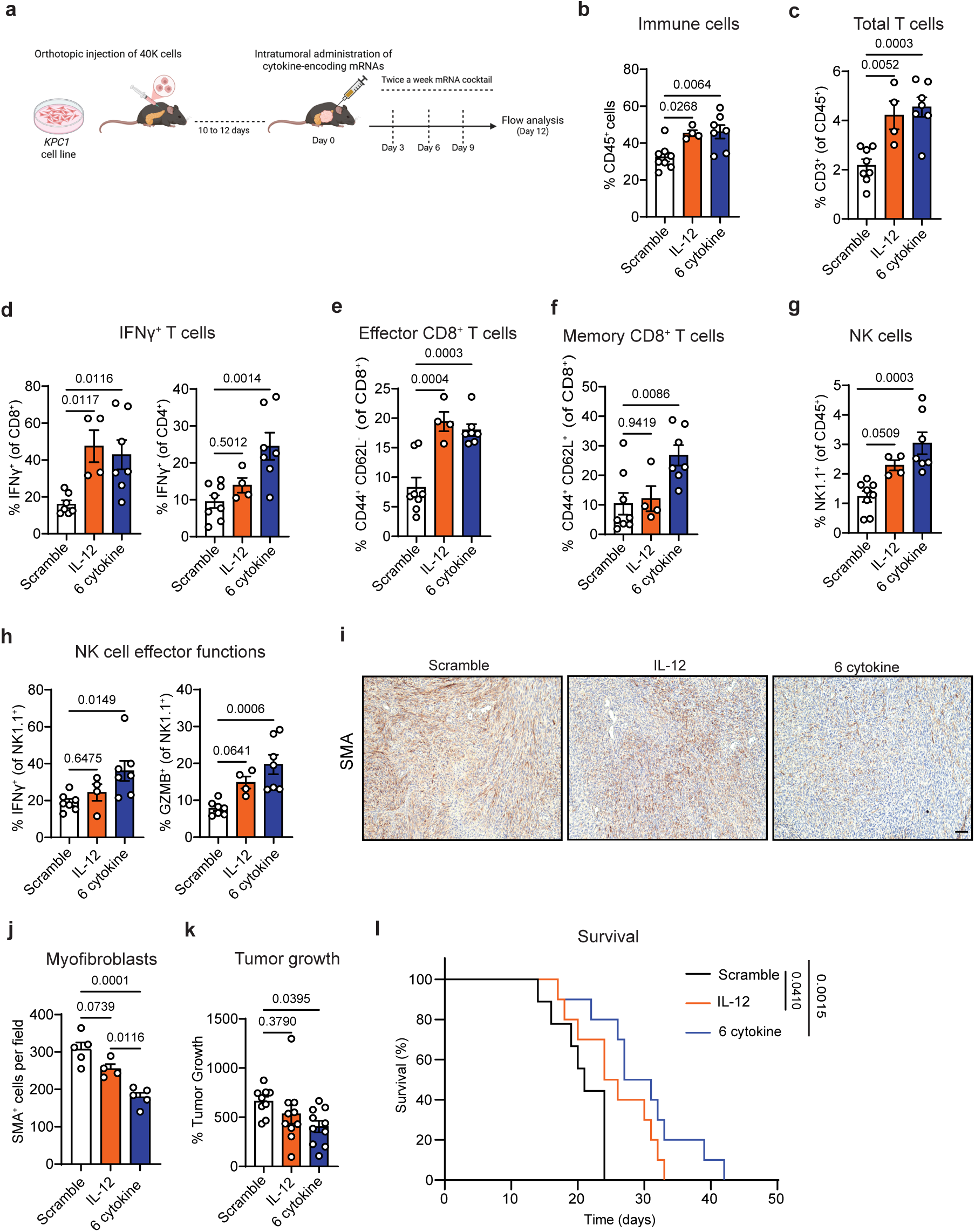
A multiplexed cytokine mRNA cocktail mobilizes innate and adaptive immunity and reduces tumor growth and desmoplasia in “cold” PDAC-bearing mice. **a**, *KPC1* PDAC tumor cells expressing luciferase-GFP were orthotopically injected into the pancreas of 8-to 12-week-old female C57BL/6 mice. Following tumor establishment, mice received intratumoral injections of either scramble mRNA (60 µg), IL-12 mRNA (10 µg), or a 6-cytokine mRNA cocktail (10 µg each of IL-12, IL-15, IL-18, IFNβ1, CCL5, and CXCL10; 60 µg total). Injections were administered every 3 days, and mice sacrificed for downstream tumor analysis on Day 12. **b-h**, Flow cytometry analysis of total CD45^+^ immune cells **(b)**, total CD3^+^ T cells **(c)**, IFNγ^+^ T cells **(d)**, effector CD8^+^ T cells **(e)**, memory CD8^+^ T cells **(f)**, total NK1.1^+^ NK cells **(g)**, and NK cell effector functions **(h)** in *KPC1* orthotopic PDAC tumors following repeated mRNA treatment as in **a** (*n* = 4–8 mice per group). **i**, Representative immunohistochemical (IHC) staining of *KPC1* orthotopic PDAC tumors from mice treated for 12 days as in **a**. Scale bar, 100 µm. **j**, Quantification of SMA^+^ myofibroblasts per field from IHC analysis in **i** (*n* = 4-5 mice per group). **k,** Ultrasound imaging analysis of the percent change in tumor volume at Day 14 as compared to Day 0 (pre-treatment baseline) in *KPC1* orthotopic tumor-bearing mice after receiving five intratumoral injections on Days 0, 3, 6, 9, and 12 with either scramble mRNA (60 µg), IL-12 mRNA (10 µg), or a 6-cytokine mRNA cocktail (10 µg each of IL-12, IL-15, IL-18, IFNβ1, CCL5, and CXCL10; 60 µg total) (*n* = 9–10 mice per group). **l**, Kaplan–Meier survival curve of mice with *KPC1* orthotopic PDAC tumors treated intratumorally with indicated mRNAs every 3 days (*n* = 9–10 mice per group). *P* values in **b-h** and **k** were calculated using ordinary one-way ANOVA with Dunnett’s correction, and those in **j** were calculated using ordinary one-way ANOVA with Tukey’s correction. *P* values in **l** were calculated using a log-rank test. Error bars are mean ± s.e.m.

Flow cytometry analysis performed after four intratumoral injections of cytokine-encoding mRNAs revealed a sustained increase in CD45^+^ immune cell infiltration into the “cold” PDAC TME in both the IL-12 mRNA alone and 6-cytokine mRNA cocktail treated groups (Fig. 3b). Total CD3^+^ T cell numbers also increased with single IL-12 or combinatorial cytokine mRNA administration, though relative proportions of CD4^+^ and CD8^+^ T cell populations were not significantly altered (Fig. 3c and Extended Data Fig. 2a). IFNγ positivity, indicative of effector cytokine production, was significantly increased in CD8^+^ T cells in both cytokine treatment arms and in CD4^+^ T cells only in mice receiving the 6 cytokine mRNA cocktail as compared to Scramble mRNA controls (Fig. 3d). The proportion of CD44^+^CD62L^-^ effector CD8^+^ T cells were also significantly augmented following single IL-12 or combined cytokine mRNA delivery (Fig. 3e). However, despite increased effector differentiation, CD8^+^ T cells were not cytotoxic following cytokine mRNA therapy in this model, as measured by Granzyme B (GZMB) positivity (Extended Data Fig. 2b).

Strikingly, the combinatorial 6 cytokine mRNA cocktail, but not single IL-12 mRNA administration, increased the percentage of memory CD8^+^ T cells in *KPC* PDAC tumors (Fig. 3f). There was also a trend toward increased numbers of MHC-II^+^ DCs following the 6 cytokine mRNA therapy that are important for priming and recall of memory T cells^47^ (Extended Data Fig 2c). Moreover, there were significant changes in the NK1.1^+^ NK cell population following combinatorial mRNA cytokine administration that could not be achieved with IL-12 mRNAs alone. Not only did NK cell numbers that are typically lacking in the PDAC TME increase, but their expression of both IFNγ and the cytotoxicity marker GZMB were also enhanced upon repeated 6 cytokine mRNA dosing (Fig. 3g,h). Additionally, while there was no change in F4/80^+^ macrophage populations, Gr-1^+^ myeloid-derived suppressor cell (MDSC) numbers were modestly increased with 6 cytokine mRNA dosing (Extended Data Fig. 2d), which could contribute to some of the failure to fully activate cytotoxic T cells. Thus, despite similar effects on effector T cell responses between both single IL-12 and 6 cytokine mRNA therapy, the combined mRNA cocktail was able to further mobilize memory CD8^+^ T cell formation and activate cytotoxic NK cell responses in this “cold” PDAC model.

Treatment with the 6 cytokine mRNA cocktail also had a marked impact on PDAC tumors and their surrounding stroma. Myofibroblasts prevalent within PDAC lesions contribute to a desmoplastic TME that can restrict the infiltration of NK and T cells^48^. Immunohistochemical (IHC) analysis revealed a reduction of smooth muscle actin (SMA)⁺ myofibroblasts following IL-12 mRNA monotherapy that was even more pronounced in tumors receiving the combinatorial mRNA cocktail (Fig. 3i,j), demonstrating cytokine mRNA therapy can also alleviate fibrosis in the PDAC TME that could indirectly impact immunity. Similarly, whereas single IL-12 mRNA therapy had no impact on tumor growth following biweekly intratumoral dosing over 14 days, the 6-mRNA cocktail achieved a significant reduction in tumor growth as assessed by ultrasound imaging (Fig. 3k). Importantly, these cytokine mRNAs had minimal impact of *KPC* tumor cell and CAF viability *in vitro*, suggesting that the anti-tumor and fibrotic effects observed *in vivo* are unlikely the result of the direct effects of cytokine production on tumor cells and fibroblasts (Extended Data Fig. 2e,f). Furthermore, long-term studies revealed that while IL-12 mRNA monotherapy marginally increased survival in mice compared to control Scramble mRNA dosing, combined 6 cytokine mRNA cocktail treatment more significantly enhanced overall survival outcomes (Fig. 3l). Together, these results demonstrate that multiplexed cytokine mRNA therapy can promote both NK and T cell effector activation and memory and produce anti-tumor responses and fibrosis resolution in PDAC.

### Combinatorial cytokine mRNA therapy can achieve robust cytotoxic T cell responses in a “hot” PDAC model expressing neo-antigens

The *KPC1* orthotopic PDAC model is immunologically “cold” and lacks neo-antigens that may be necessary for effective cytotoxic T cell responses^12^. To test our cytokine mRNA cocktail in a “hot” PDAC model, we orthotopically implanted the syngeneic *KPCY 2838c3* cell line, which exhibits a T cell-high phenotype and is characterized by the presence of three predicted neoantigens^49^, into C57BL/6 mice. Based on our earlier *in vitro* findings that IL-15 was not necessary for inducing NK and CD8^+^ T cell migration and effector functions with respect to the 6 cytokine combination (see Fig. 2a-f), we moved forward with a refined 5 cytokine mRNA cocktail (IL-12, IL-18, CCL5, CXCL10, IFNβ) that excluded IL-15 for subsequent studies.

Following *KPCY 2838c3* tumor induction (as assessed by ultrasound), control Scramble mRNA, IL-12 mRNA alone, or the 5 cytokine mRNA combination was injected intratumorally twice per week (Fig. 4a). Flow cytometry analysis after 12 day treatment showed an increase in total CD45⁺ immune cell infiltration accompanied by a significant expansion of both total as well as CD8⁺ and CD4⁺ T cell subsets in the 5 cytokine mRNA cocktail but not single IL-12 mRNA group (Fig. 4b-d). Though both IL-12 alone and combined cytokine mRNAs enhanced the percentage of effector and memory CD8^+^ T cells in the PDAC TME compared to control Scramble mRNA, a trend toward increased IFNγ and GZMB positivity was only observed in the 5 cytokine mRNA arm (Fig. 4e and Extended Data Fig. 3a). CD4^+^ T cell effector functions were unaltered by cytokine mRNA administration in this model (Extended Data Fig. 3b). Despite no change in total NK cell numbers in this T cell-high *KPCY* PDAC model, there was a significant increase in their expression of effector cytokine IFNγ and cytotoxic GZMB granules following cytokine therapy (Fig. 4f and Extended Data Fig. 3c). There was also a significant increase in CD103a^+^ cross-presenting DCs and MHC-II^+^ macrophages, as well as a decrease in MDSCs in both single and combinatorial cytokine mRNA conditions that could further support cytotoxic lymphocyte priming and activity in the PDAC TME (Fig. 4g,h). Ultrasound imaging of PDAC tumors revealed that while repeated IL-12 mRNA dosing alone did not alter tumor growth, treatment with the 5 cytokine mRNA cocktail led to significant tumor growth reduction after 12 day treatment compared to Scramble mRNA (Fig. 4i). Thus, the 5 cytokine mRNA combination outperforms IL-12 single mRNA intratumoral injections in its ability to elicit CD8^+^ T cell infiltration and effector activation and tumor growth control in a “hot” PDAC model.

**Fig 4.**
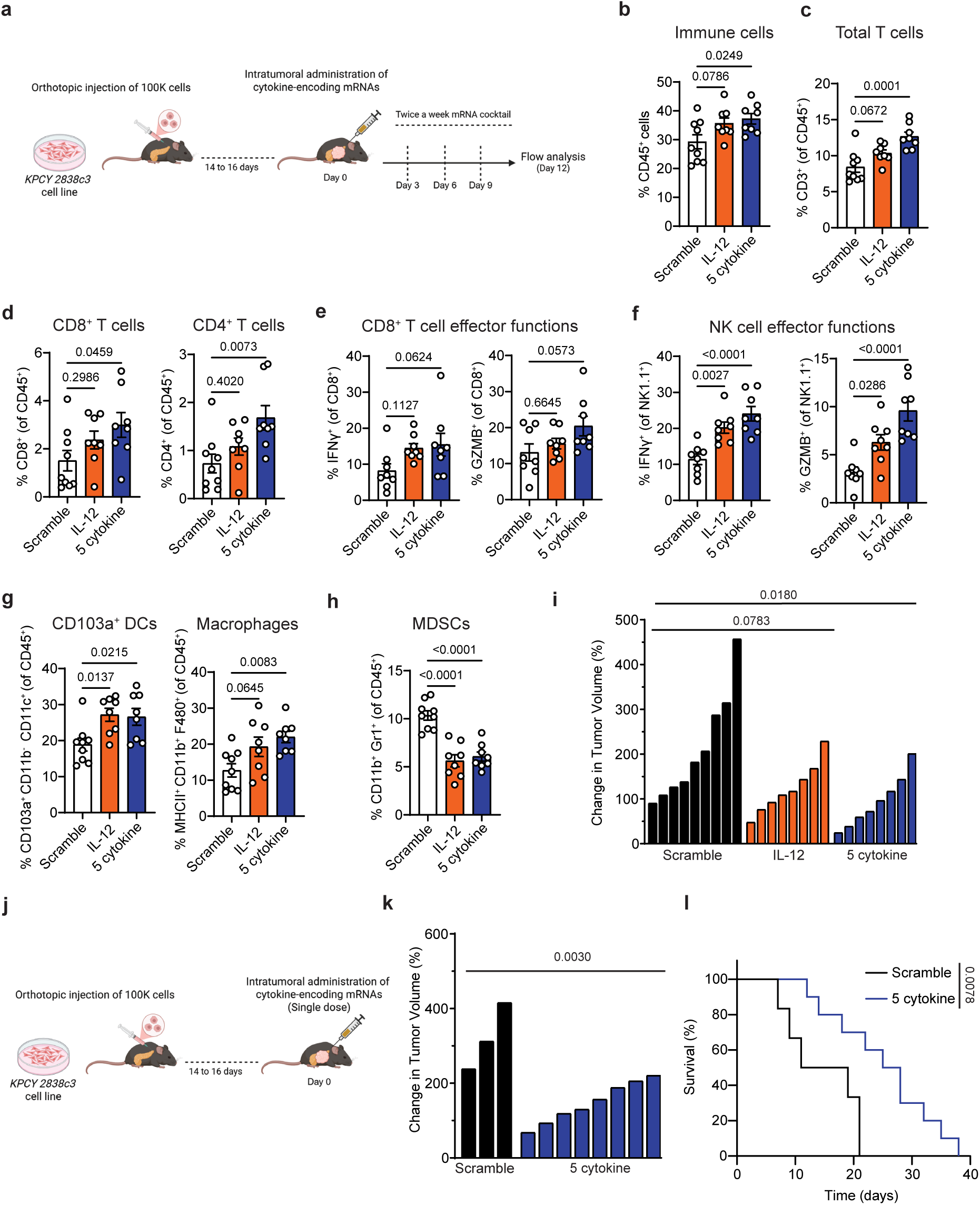
Cytokine mRNA therapy can activate cytotoxic T cell immunity in a “hot” PDAC model. **a,** *KPCY 2838c3* PDAC tumor cells were orthotopically injected into the pancreas of 8-to 12-week-old female C57BL/6 mice. Following tumor establishment, mice received intratumoral injections of either scramble mRNA (50 µg), IL-12 mRNA (10 µg), or a 5-cytokine mRNA cocktail (10 µg each of IL-12, IL-18, IFNβ1, CCL5, and CXCL10; 50 µg total). Injections were administered every 3 days, and mice were sacrificed for downstream tumor analysis on Day 12. **b-h**, Flow cytometry analysis of CD45^+^ immune cells **(b)**, total CD3^+^ T cells **(c)**, CD8^+^ and CD4^+^ T cells **(d)**, CD8^+^ T cell effector functions **(e)**, NK cell effector functions **(f)**, CD103a^+^ cross-presenting dendritic cells (DCs) and MHC-II^+^ macrophages **(g)**, and Gr-1^+^ myeloid-derived suppressor cells (MDSCs) **(h)** in *KPCY 2838c3* orthotopic PDAC tumors following indicated mRNA treatment as in **a** (*n* = 8–9 mice per group). **i**, Waterfall plot of the response of *KPCY 2838c3* orthotopic PDAC tumors to treatment as in **a** (*n* = 8–9 mice per group). **j**, *KPCY 2838c3* PDAC tumor cells were orthotopically injected into the pancreas of 8-to 12-week-old female C57BL/6 mice. Following tumor establishment, mice received a single intratumoral injection of either scramble mRNA (50 µg) or a 5-cytokine mRNA cocktail (10 µg each of IL-12, IL-18, IFNβ1, CCL5, and CXCL10; 50 µg total), and tumor growth and survival were monitored. **k**, Waterfall plot of the response of *KPCY 2838c3* orthotopic PDAC tumors to a single intratumoral dose of mRNAs as in **j** (*n* = 3–8 mice per group). **l**, Kaplan–Meier survival curve of mice with *KPCY 2838c3* orthotopic PDAC tumors treated with a single intratumoral injection of mRNAs as in **j** (*n* = 6-10 mice per group). *P* values in **b-i** were calculated using ordinary one-way ANOVA with Dunnett’s correction, and those in **k** using two-tailed, unpaired Student’s t-test. *P* values in **l** were calculated using a log-rank test. Error bars are mean ± s.e.m.

Given the robustness of these innate and adaptive immune responses to repeated cytokine mRNA administration, we were curious if a single injection of the 5 cytokine mRNA cocktail could achieve similar outcomes (Fig. 4j). Remarkably, 2 weeks after a single intratumoral mRNA injection, ultrasound imaging revealed significant tumor growth control in the 5-cytokine mRNA group compared to the cohort receiving control Scramble mRNA (Fig. 4k). Even more strikingly, a single dose of the 5 cytokine mRNAs was able to significantly increase overall survival in mice bearing *KPCY 2838c3* PDAC tumors (Fig. 4l). Overall, these anti-tumor phenotypes produced in an immunologically “hot” PDAC model, even after a single mRNA dose, suggest that the presence of tumor antigens may further enhance the robustness of cytokine therapies to fully activate a coordinated innate and adaptive immune response to sustain T cell immunity against PDAC.

### Co-delivery of cytokine- and antigen-encoding mRNAs elicits a comprehensive anti-tumor immune response in a “cold” PDAC model

Our above findings highlight the synergistic potential of combining cytokine-encoding mRNAs with strategies that could enhance tumor-specific T cell priming in PDAC. Recent clinical trials have explored neo-antigen-encoding mRNA vaccines as a therapeutic strategy for PDAC, demonstrating improved patient survival through the induction of long-lasting tumor antigen-specific T cell immunity^50,51^. Hence, we hypothesized that adding mRNAs encoding tumor-associated antigens (TAAs) to our existing cytokine mRNA cocktail as an all-in-one therapy approach could further enhance and prolong anti-tumor T cell responses, particularly in the setting of a “cold” PDAC TME.

To this end, we generated mRNAs encoding three well-characterized PDAC TAAs—Mesothelin (MSLN)^52^, Mucin-1 (MUC-1)^53^, and Prostate-Specific Membrane Antigen (PSMA)^54^. Upon intratumoral delivery of these 3 antigen-encoding mRNAs as a cocktail into the *KPC1* PDAC tumor model, MSLN, MUC-1, and PSMA protein surface expression were measurably increased in tumors 24 hours post-injection as compared to control Scramble mRNA injection as assessed by IHC analysis (Fig. 5a). To determine if intratumoral delivery of these TAA mRNAs could effectively generate antigen-specific T cells within the PDAC TME, we devised an *ex vivo* antigen stimulation assay. Splenocytes were first isolated from wild-type C57BL/6 mice and pulsed with antigen-encoding mRNAs for 24 hours *ex vivo*. In parallel, T cells were isolated from tumors of PDAC-bearing mice that had been vaccinated intratumorally with the same antigen-encoding mRNAs 72 hours prior (Extended Data Fig. 4a). Antigen-specific activation of CD8^+^ T cells from tumor-bearing mice vaccinated with the 3 antigen mRNA cocktail was confirmed by a marked increase in IFNγ⁺ GZMB⁺ CD8⁺ T cells following exposure to antigen-mRNA-pulsed splenocytes (Extended Data Fig. 4b).

**Fig. 5.**
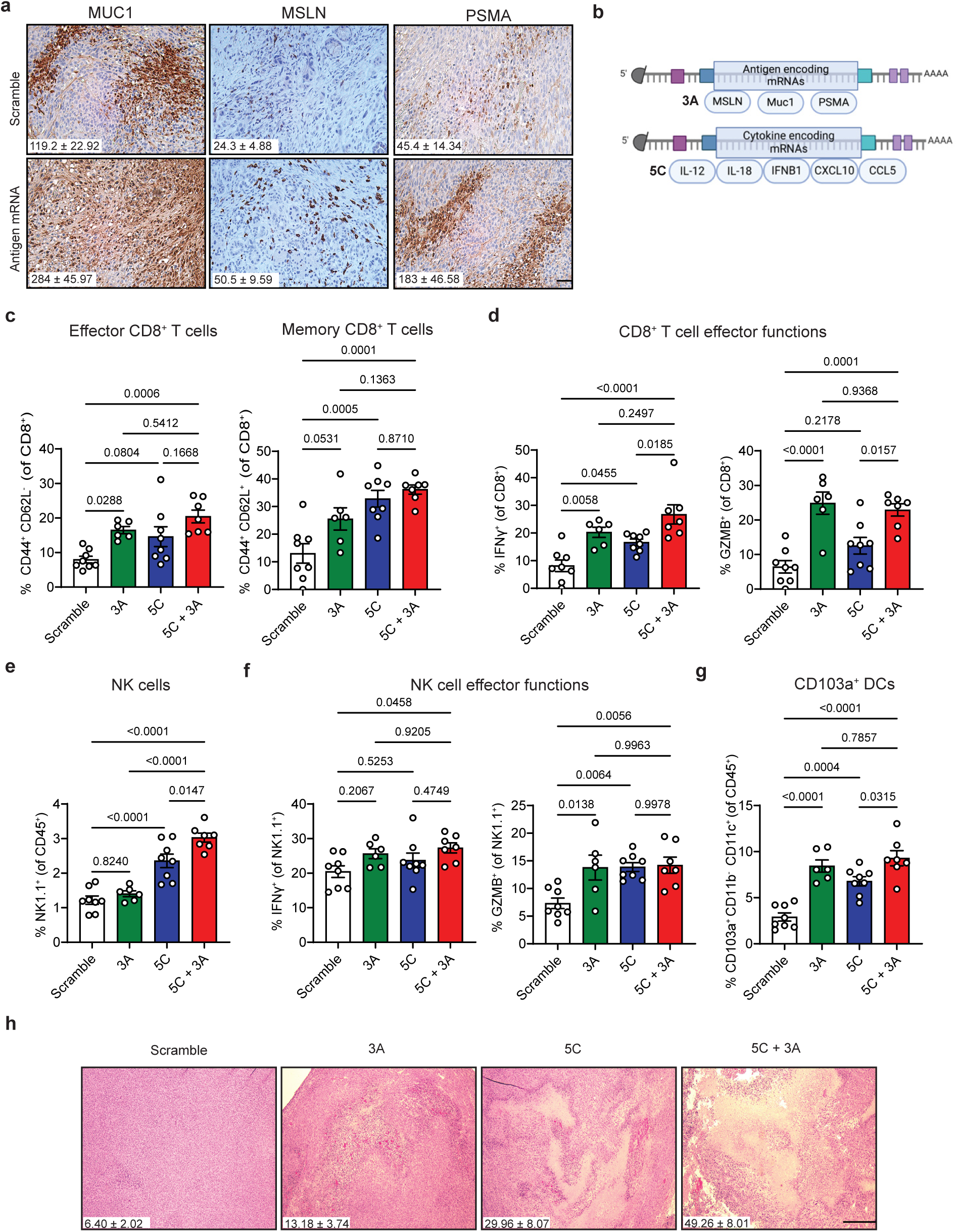
Combined cytokine- and antigen-encoding mRNA delivery potentiates anti-tumor immunity in “cold” PDAC model. **a**, Representative IHC staining of *KPC1* orthotopic PDAC tumors from mice 24 hours after intratumoral injection of either scramble mRNA (30 µg) or a 3-antigen mRNA cocktail (10 µg each of MUC-1, MSLN, and PSMA; 30 µg total). Scale bar, 100 μm. Quantification of the number of MUC-1^+^, MSLN^+^, and PSMA^+^ cells per field is shown in the inset (*n* = 5 mice per group). **b**, Schematic of the 3-antigen (3A) (MUC-1, MSLN, PSMA) and 5-cytokine (5C) (IL-12, IL-18, IFNβ1, CCL5, CXCL10) mRNA cocktails. **c-g**, Flow cytometry analysis of effector and memory CD8^+^ T cells **(c)**, CD8^+^ T cell effector functions **(d)**, NK1.1^+^ NK cells **(e)**, NK cell effector functions **(f)**, and CD103a^+^ cross-presenting DCs **(g)** in *KPC1* orthotopic PDAC tumors on Day 12 following intratumoral injections of either scramble mRNA (80 µg), the 3A mRNA cocktail (10 µg each of MUC-1, MSLN, and PSMA; 30 µg total), the 5C mRNA cocktail (10 µg each of IL-12, IL-18, IFNβ1, CCL5, and CXCL10; 50 µg total), or both 3A and 5C in combination every 3 days (*n* = 6-8 mice per group). **h**, H&E staining of *KPC1* orthotopic PDAC tumors following 12 day treatment of mice as in **c**. Scale bar, 500μm. Quantification of the percentage of tumor area covered in necrosis is shown in the inset (*n* = 6-8 mice per group). *P* values in **c-g** were calculated using ordinary one-way ANOVA with Tukey’s correction. Error bars are mean ± s.e.m.

Having validated the capacity of the 3 antigen mRNA combination to generate antigen-specific T cells in PDAC tumors, we set out to investigate the impact of these engineered TAA mRNAs alone or as part of an all-in-one cocktail with the 5 cytokine-encoding mRNA combination on immune responses in the cold *KPC1* PDAC mouse model (Fig. 5b). Though repeated intratumoral administration of the 3 antigen-encoding mRNAs (3A) alone or in combination with the 5 cytokine-encoding mRNAs (5C + 3A) had no effect on total immune cell frequencies or percentages of CD8⁺ or CD4⁺ T cells in the PDAC TME after 12 days of treatment (Extended Data Fig. 4c,d), there was a significant increase in the percentage of effector and memory CD8⁺ T cells in the PDAC TME compared to Scramble controls (Fig. 5c). Antigen mRNA dosing was also sufficient to enhance GZMB and IFNγ expression in CD8^+^ T cells, and in combination with cytokine mRNAs significantly increase CD8^+^ T cell IFNγ and GZMB positivity compared to the 5 cytokine mRNA cocktail alone (Fig. 5d). Combined antigen and cytokine mRNA administration also significantly enhanced IFNγ production from NK cells and CD4^+^ T cells and numbers of antigen-presenting macrophages compared to Scramble controls, and even further increased NK cell and CD103a^+^ DC accumulation compared to 5 cytokine mRNA therapy alone (Fig. 5e-g and Extended Data Fig. 4e-f). MDSC numbers, on the other hand, remained unchanged across conditions (Extended Data Fig. 4g).

This combined effect on DC antigen presentation and NK and CD8^+^ T cell activity was associated with extensive tumor necrosis, as visible by gross histology, in 5C + 3A treated mice as compared to the single antigen or cytokine mRNA monotherapy arms (Fig. 5h). Ultrasound analysis further demonstrated that dosing with the combination of 3 antigen and 5 cytokine-encoding mRNAs resulted in significant tumor growth suppression in *KPC1* PDAC transplant mice (Extended Data Fig. 4h). Collectively, these results highlight the synergistic potential of combining cytokine and antigen mRNA therapy modalities to overcome the cold PDAC TME and produce anti-tumor immune responses.

### A single dose of a multiplexed antigen and cytokine mRNA cocktail is sufficient to confer NK and CD8^+^ T cell-mediated PDAC control

As repeated intratumoral mRNA therapy administration may not be feasible in patients with pancreatic cancer in the clinic, we set out to determine if a single dose of an all-in-one cocktail containing 5 cytokine (IL-12, IL-18, CCL5, CXCL10, IFNβ) and 3 antigen (MSLN, MUC-1, PSMA) mRNAs would be sufficient to produce durable immune-mediated anti-tumor effects in our PDAC animal models (Fig. 6a). A single intratumoral dose of the combined cytokine and antigen mRNA cocktail in established “cold” *KPC1* PDAC tumors led to a significant decrease in tumor growth 2-weeks post-treatment that culminated in prolonged overall survival outcomes compared to mice treated with control Scramble mRNAs (Fig. 6b,c). Though a single dose of the 5 cytokine mRNA mix on its own was also able to confer a survival benefit in PDAC-bearing animals, this survival advantage was significantly prolonged when combined with the 3 antigen mRNAs (Fig. 6d).

**Fig. 6.**
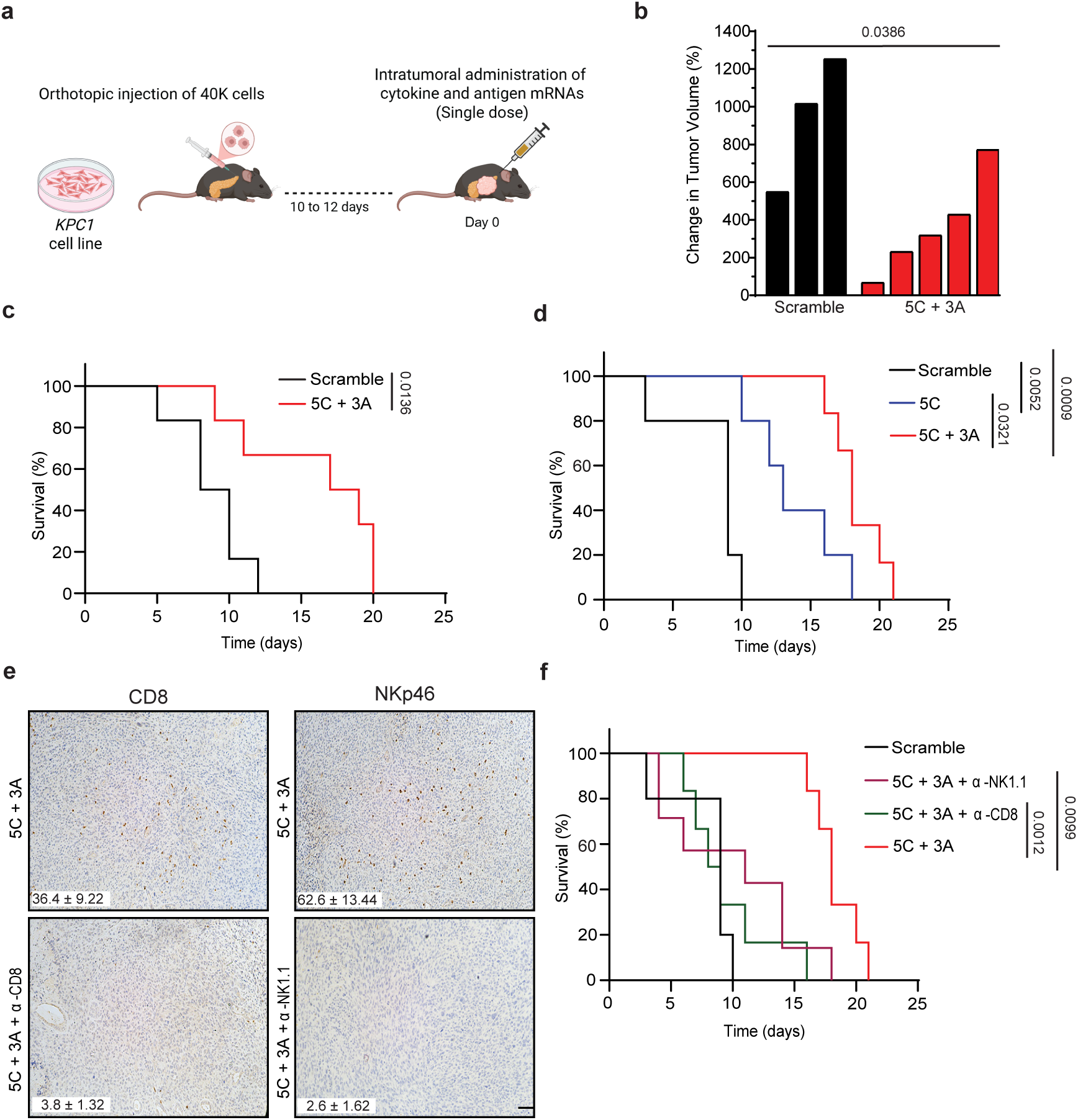
A single dose of an all-in-one antigen and cytokine mRNA cocktail is sufficient for NK and CD8^+^ T cell-mediated PDAC control. **a**, *KPC1* PDAC tumor cells expressing luciferase-GFP were orthotopically injected into the pancreas of 8-to 12-week-old female C57BL/6 mice. Following tumor establishment, mice received a single intratumoral injection of either scramble mRNA (80 µg) or the combined 5-cytokine (5C) and 3-antigen (3A) mRNA cocktail (10 µg each of IL-12, IL-18, IFNβ1, CCL5, CXCL10, MUC-1, MSLN, and PSMA; 80 µg total), and tumor growth and survival were monitored. **b**, Waterfall plot of the response of *KPC1* orthotopic PDAC tumors to single dose mRNA therapy as in **a** (*n* = 3-5 mice per group). **c**, Kaplan–Meier survival curve of mice with *KPC1* orthotopic PDAC tumors treated intratumorally with single dose mRNA therapy in **a** (*n* = 6 mice per group). **d**, Kaplan–Meier survival curve of mice with *KPC1* orthotopic PDAC tumors treated intratumorally with a single dose of scramble mRNA (80 µg), the 5-cytokine mRNA cocktail (10 µg each of IL-12, IL-18, IFNβ1, CCL5, and CXCL10; 50 µg total), or the combined 5-cytokine and 3-antigen mRNA cocktail (10 µg each of IL-12, IL-18, IFNβ1, CCL5, CXCL10, MUC-1, MSLN, and PSMA; 80 µg total) (*n* = 5-6 mice per group). **e**, Representative IHC staining of *KPC1* orthotopic PDAC tumors harvested at endpoint from mice treated with a single intratumoral injection of a combined 5-cytokine and 3-antigen mRNA cocktail (10 µg each of IL-12, IL-18, IFNβ1, CCL5, CXCL10, MUC-1, MSLN, and PSMA; 80 µg total) alone or in combination with NK1.1 (PK136; 250 µg) or CD8 (2.43; 200 µg) depleting antibodies administered twice per week following tumor formation. Scale bar, 100 μm. Quantification of the number of CD8^+^ and NKp46^+^ NK cells per field is shown in the inset (*n* = 6-7 mice per group). **f,** Kaplan–Meier survival curve of mice with *KPC1* orthotopic PDAC tumors treated intratumorally with a single dose of scramble mRNA or the combined 5C and 3A cocktail in the presence or absence of NK1.1 (PK136) or CD8 (2.43) depleting antibodies as in **e** (*n* = 5-7 mice per group). *P* values in **b** were calculated using two-tailed, unpaired Student’s t-test, and those in **c-d** and **f** were calculated using a log-rank test.

To determine whether the observed anti-tumor effects of mRNA therapy were indeed immune-dependent, some mice that received the combined 3 antigen and 5 cytokine mRNA cocktail injection were also administered NK1.1 (PK136) and CD8 (2.43) monoclonal antibodies simultaneously to deplete NK cell and CD8^+^ T cell populations, respectively (Fig. 6e). Both NK and CD8⁺ T cell ablation abrogated the survival benefit conferred by combinatorial cytokine and antigen mRNA therapy (Fig. 6f). Thus, a single intratumoral dose of an immunogenic mRNA cocktail is sufficient to orchestrate cytotoxic lymphocyte-mediated tumor suppression and extend survival outcomes.

### NP-mediated delivery of cytokine and antigen mRNAs achieves complete tumor responses in an autochthonous PDAC mouse model

To further evaluate the therapeutic potential of our immunogenic cytokine- and antigen-encoding mRNA cocktail in a more clinically relevant setting, we expanded our studies into *KPC* genetically engineered mouse models (GEMMs) of PDAC. Given that GEMMs develop spontaneous and multi-focal tumors throughout the pancreas that generally preclude direct intratumoral injection, we encapsulated mRNAs into pegylated lipid nanoparticles (NPs) to enable systemic delivery via tail vein injection. We successfully encapsulated the 5 cytokine mRNAs into NPs using ionizable lipids at high efficiency to produce an mRNA concentration (∼10 µg per mRNA) comparable to our direct injection of naked mRNAs (Extended data Fig. 5a,b). These mRNA-loaded NPs also had a neutral charge and a small 50-60 nM diameter (Extended Data Fig. 5c,d), which we have previously shown to enable homing to and retention of NPs carrying immune agonists in the PDAC TME following systemic delivery with minimal toxicity and off-target inflammatory effects^15^.

Indeed, while there was a transient spike in cytokine levels in the blood and liver 24 hours following intravenous (i.v.) delivery of NPs loaded with the 5 cytokine mRNA cocktail into PDAC-bearing *KPC* GEMM mice, cytokine levels were mostly restored to baseline (with the exception of IFNβ in the liver and IL-18 in the blood) 48 hours post-administration (Fig. 7a and Extended Data Fig. 5e,f). In contrast, the protein quantities of IL-12, IL-18, IFNβ, CCL5, and CXCL10 remained significantly elevated within PDAC tumors 48 hours after i.v. injection of NPs encapsulated with the 5 cytokine mRNA cocktail as compared to control Scramble mRNAs (Fig. 7b). Importantly, i.v. administration of 5 cytokine mRNA-encapsulated NPs led to no observable systemic toxicities as measured by changes in animal body weight (Extended Data Fig. 5g). Systemic administration of NPs loaded with the 3 antigen mRNA cocktail also led to increased protein expression of MUC-1, MSLN, and PSMA in *KPC* GEMM PDAC lesions 48 hrs later as assessed by IHC analysis (Fig. 7c). Thus, systemic administration of mRNAs through NP formulations is a safe and effective strategy to enhance cytokine and antigen expression in PDAC tumors in autochthonous GEMMs.

**Fig. 7.**
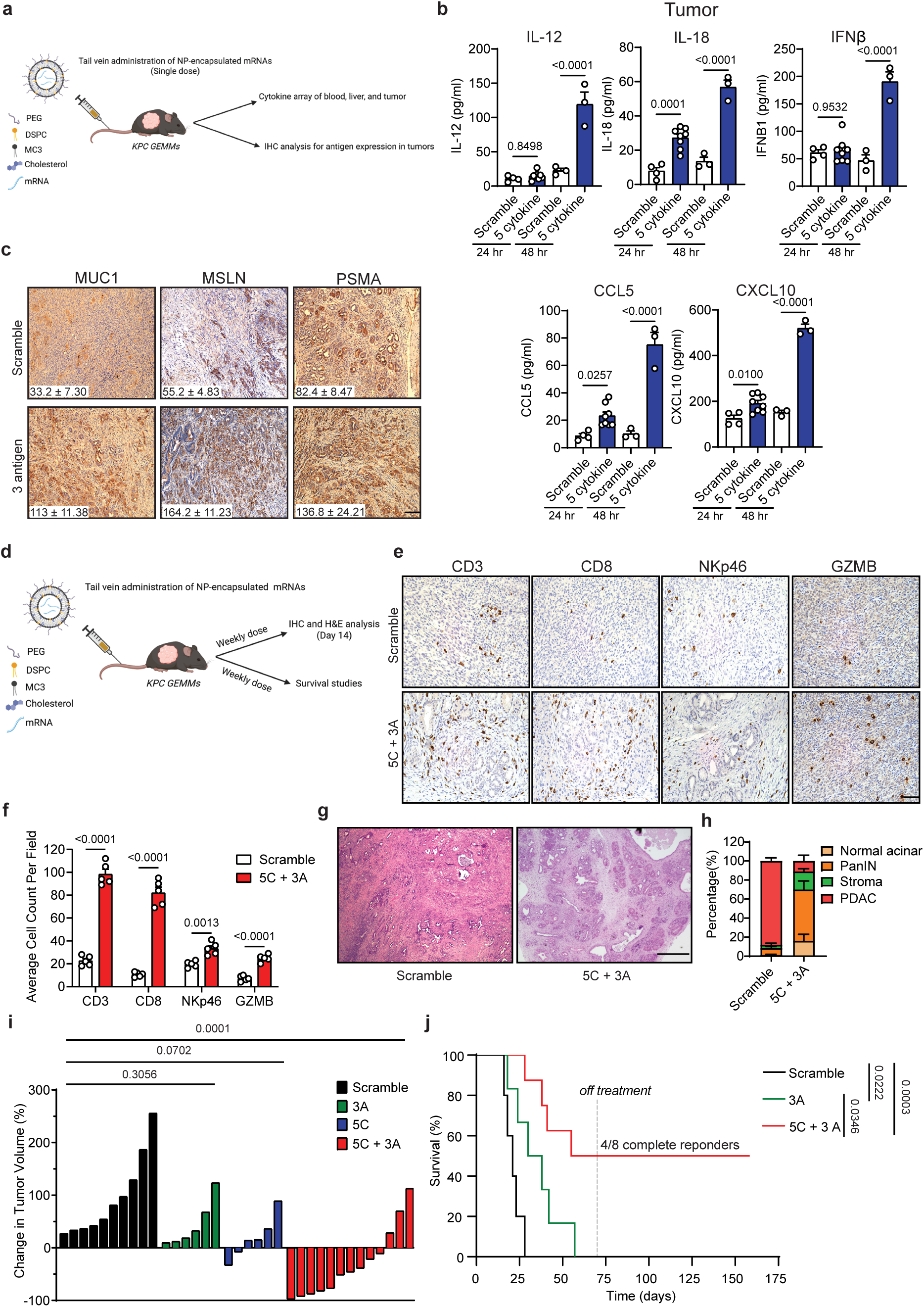
Systemic delivery of NP-encapsulated cytokine and antigen mRNAs produces complete tumor responses and long-term survival in autochthonous PDAC GEMMs. **a**, *KPC* GEMMs with confirmed tumors received a tail vein injection of lipid nanoparticles (NPs) encapsulated with scramble mRNA (50 µg), the 5-cytokine mRNA cocktail (10 µg each of IL-12, IL-18, IFNβ1, CCL5, and CXCL10; 50 µg total), or the 3-antigen mRNA cocktail (10 µg each of MUC-1, MSLN, and PSMA; 30 µg total). Tumor, liver, and blood samples were collected at 24- and 48-hours post-injection for cytokine array and IHC analysis. **b**, Cytokine array results from tumor tissues collected at different time points following systemic administration of scramble or 5 cytokine mRNA NPs as in **a** (*n* = 3–8 mice per group). **c**, Representative IHC staining of *KPC* GEMM tumors collected 48 hours post-injection of NPs encapsulated with scramble mRNA (30 µg) or the 3-antigen mRNA cocktail (10 µg each of MUC-1, MSLN, and PSMA; 30 µg total). Scale bar, 100 μm. Quantification of the number of MUC-1^+^, MSLN^+^, and PSMA^+^ cells per field is shown in the inset (*n* = 5 mice per group). **d**, *KPC* GEMMs with confirmed tumors received weekly tail vein injections of NPs encapsulated with scramble mRNA (80 µg), the 5-cytokine (5C) mRNA cocktail (10 µg each of IL-12, IL-18, IFNβ1, CCL5, and CXCL10; 50 µg total), the 3-antigen (3A) mRNA cocktail (10 µg each of MUC-1, MSLN, and PSMA; 30 µg total), or combined 5C and 3A mRNA cocktails (80 µg total). Mice were either sacrificed at Day 14 for IHC and H&E analysis or treated until end-point for long-term survival analysis. **e**, Representative IHC staining of *KPC* GEMM tumors 14 days after weekly mRNA NP dosing as in **d**. Scale bar, 100 μm. **f**, Quantification of the number of CD3^+^ T cells, CD8^+^ T cells, NKp46^+^ NK cells, and GZMB^+^ cytotoxic cells per field from IHC analysis in **e** (*n* = 5 mice per group). **g**, Representative H&E staining of *KPC* GEMM tumors 14 days after weekly mRNA NP dosing as in **d**. Scale bar, 500 μm. **h**, Quantification of the percentage of normal acinar, stromal, pancreatic intraepithelial neoplasia (PanIN), and adenocarcinoma pathology in pancreas tumors from *KPC* GEMM animals treated for 14 days as in **d** (*n* = 4-5 mice per group). **i**, Waterfall plot of the response of *KPC* GEMM tumors to 14 day mRNA NP treatment as in **d** (*n* = 6-13 mice per group). **j**, Kaplan–Meier survival curve of tumor-bearing *KPC* GEMM animals treated weekly with mRNA NPs as in **d** (*n* = 5-8 mice per group). Dotted line indicates when mice were taken off treatment at Day 70. *P* values in **b** were calculated using ordinary one-way ANOVA with Sidák’s correction, and those in **f** using two-way ANOVA with Sidák’s correction. *P* values in **i** were calculated using ordinary one-way ANOVA with Dunnett’s correction, and those in **j** using a log-rank test. Error bars are mean ± s.e.m.

To assess the impact of NP-mediated cytokine and antigen mRNA delivery on immune and tumor responses in *KPC* GEMM tumors, we administrated NPs containing the all-in-one 5 cytokine and 3 antigen (5C + 3A) mRNA cocktail or Scramble mRNA controls by weekly i.v. injection following ultrasound confirmation of tumor formation (Fig. 7d). Immunophenotyping after 2-week treatment revealed a significant accumulation of total CD3^+^ and CD8^+^ T cells, NKp46^+^ NK cells, and cytotoxic GZMB^+^ immune cells in the PDAC TME of *KPC* GEMM animals treated with combined 5C + 3A NPs (Fig. 7e,f). Further histopathological examination uncovered a striking ablation of adenocarcinoma pathology, with only rare tumor foci evident amongst predominantly normal acinar structures, pre-malignant pancreatic intraepithelial neoplasia (PanIN) lesions, and stroma in the pancreas of PDAC-bearing *KPC* mice in the 5C + 3A NP treated cohort (Fig. 7g,h, and Extended Data Fig. 5h).

Combined cytokine and antigen mRNA NP delivery mediated long-term, durable pancreatic tumor control in *KPC* GEMMs that could not be achieved through NP encapsulation of antigen or cytokine mRNAs individually. Ultrasound imaging analysis showed that although antigen mRNA NP or cytokine mRNA NP dosing alone slowed tumor growth compared to the control Scramble mRNA group, there was no significant change in overall tumor burden after 2-week treatment (Fig. 7i). In contrast, treatment of PDAC-bearing *KPC* GEMMs with the combinatorial 5C + 3A mRNA NP cocktail led to significant tumor regressions in the majority in mice (10/13), which we rarely observed in 3A NP (0/6) or 5C NP (2/6) groups. Impressively, many of the PDAC tumors in mice treated with the all-in-one cytokine and antigen mRNA NPs continued to regress and even achieved complete responses as early as 8 weeks after repeated weekly NP dosing (Extended Data Fig. 5i). While PDAC-bearing *KPC* GEMM mice receiving control Scramble mRNA NPs quickly succumbed to the disease in a matter of weeks (median survival = 21 days), treatment with all-in-one 5C + 3A mRNA NPs dramatically extended overall survival by a number of months (median survival = 106.5 days) (Fig. 7j). Remarkably, 50% of the mice receiving 5C + 3A mRNA NPs had complete tumor responses that were durable for months after treatment ceased. While systemic administration of NPs encapsulated with only antigen mRNAs also led to a modest but statistically significant increase in overall survival (median survival = 34 days) compared to Scramble mRNA dosing, antigens alone were not able to produce complete responses or long-term survival benefits in *KPC* GEMM animals with established PDAC tumors (Fig. 7j). Collectively, these results demonstrate that systemic delivery of a multiplexed cytokine and antigen mRNA therapy can achieve complete and durable tumor responses in autochthonous preclinical PDAC models.

## DISCUSSION

The PDAC TME is known to contribute to immune suppression and poor drug delivery, and as such innovative strategies are needed to overcome these obstacles for effective immunotherapy^55^. Our previous work showed that therapies targeting the central KRAS signaling node in PDAC can remodel this immune suppressive TME through induction of the senescence-associated secretory phenotype (SASP), a collection of pleiotropic factors that include a diverse array of pro-inflammatory cytokines that are necessary to activate anti-tumor immunity and even potentiate immune checkpoint blockade (ICB) therapies^12,14,15^. However, senescent cell persistence and chronic SASP production can also conversely contribute to immune suppression and tumor progression through IL-6, IL-8, and other factors^56–59^. As such, we hypothesized that delivering only the cytokine signals we have shown to activate productive innate and adaptive immunity through the SASP, while bypassing the need for cellular senescence, could make an effective immunotherapy for PDAC. To this end, we utilized mRNA technology to produce transient yet robust cytokine expression in the local PDAC TME through intratumoral delivery of SASP-related cytokine mRNAs. Given the modularity of mRNA synthesis, we were able to test different combinations of interleukins, chemokines, and interferons to fine tune the optimal combination to elicit DC, NK, and CD8^+^ T cell mobilization and activation. A 5 cytokine cocktail consisting of IL-12, IL-18, CCL5, CXCL10, and IFNβ was sufficient to activate innate immunity and enhance T cell memory formation in “cold” PDAC mouse models. However, cytokine mRNAs were only able to activate CD8^+^ T cell effector functions and cytotoxicity in “hot” PDAC models containing putative neo-antigens. Combining the 5 cytokine mRNA cocktail with mRNAs encoding 3 known tumor-associated antigens (TAAs) in PDAC as an all-in-one package was sufficient to prolong survival after just a single intratumoral dose, as well as produce complete tumor responses following weekly systemic administration of NPs encapsulating these mRNAs in preclinical PDAC-bearing GEMMs by mobilizing both NK and CD8^+^ T cell anti-tumor immunity. Thus, this multiplexed, immunogenic mRNA approach is capable of reversing immune suppression and bypassing drug delivery issues to pave a path for an effective immunotherapy for PDAC.

Despite being one of the first FDA-approved immunotherapy modalities for cancer^18–21^, cytokine therapies have traditionally relied on systemic delivery of single cytokine recombinant proteins that suffer from inflammatory toxicities and modest efficacy^22–26^. The ability to now synthesize cytokine-encoding mRNAs *de novo* with nucleoside modifications to enhance stability while reducing off-target inflammatory toxicities offers a means to safely induce multiple cytokines in the TME simultaneously. Indeed, intratumoral delivery of cytokine mRNA therapies can produce immune-mediated tumor regressions in “hot” melanoma models^27–29^, and have even started to show promise in melanoma patients in early phase trials where they are well-tolerated (NCT06249048)^60^. However, whether cytokine mRNA therapies could also be effective in “cold” and immune suppressed tumors such as PDAC that are not responsive to conventional immunotherapies is unclear. A recent study showed that IL-12 mRNA could be delivered intratumorally into orthotopic PDAC lesions in mice, leading to remodeling of myeloid-mediated immune suppression and activating T cell proliferation and effector functions^30^. Still, IL-12 mRNA therapy was not sufficient, and required additional stereotactic body radiation therapy in combination, in order to fully activate anti-tumor T cell responses in PDAC models. Our work builds upon these findings to show that while IL-12 mRNAs alone can increase T cell infiltration and effector differentiation, combining IL-12 with mRNAs encoding other interleukins (IL-18), chemokines (CCL5, CXCL10), and interferon β is necessary to produce T cell activation and memory formation, as well as drive the accumulation and activation of NK cells. While previous mRNA cocktail iterations tested in other preclinical models or in cancer patients have included ILs and IFNs, we are the first to combine these inflammatory signals with chemokines, which we have previously shown are key to promoting CD8 and NK cell chemotaxis into the “cold” TME as part of the SASP^14^. Moreover, we found that delivery of these 5 cytokine encoding mRNAs leads to later induction of other cytokines, such as GM-CSF important for DC activity, as well as chemokines CCL3, CCL4, and CXCL10 important for DC, NK, and T cell migration, which could further amplify the immune response. In the future, antibody-mediated depletion of delivered or induced cytokines could be used to tease out their relative contributions to anti-tumor immunity and to further optimize our mRNA cocktail for PDAC.

While our 5 cytokine mRNA therapy induces innate and adaptive immune infiltration and NK cell activation in the TME of “cold” PDAC models, it is not sufficient to fully activate antigen-dependent T cell responses. By combining cytokine mRNAs with those encoding known TAAs, we are able to fully activate cytotoxic T cell responses in “cold” transplant and immune excluded GEMM PDAC-bearing animals. Interestingly, we found that both intratumoral delivery of naked mRNAs, as well as i.v. delivery of NP-encapsulated mRNAs, could induce similar induction of cytokine and antigen levels and immune activation. As antigen vaccines are typically delivered via intramuscular or subcutaneous injection to enhance DC uptake in the lymph nodes and systemic mobilization of T cell responses^61,62^, it would be of interest to compare other delivery routes for our mRNA therapies. mRNAs themselves could also be further modified to enhance stability and protein production through use of circularized^63,64^ or self-amplifying^65^ RNAs, as well as formulated in hydrogels^66,67^ to facilitate their slow release and retention in TME. Similarly, NPs could be engineered with targeting peptides to not only increase their uptake by tumor cells but possibly direct them to different cell types of interest in the TME to further increase potency while reducing systemic off-target effects. The modularity of our mRNA platform could allow us to multiplex other cytokines and antigens of interest, such as FLT3L^68^ to expand DCs, IL-7 to further enhance T cell memory^69^, or other putative TAAs in PDAC (e.g. EGFR^70^, CLDN18.2^71^, MUC16^72^) to target additional heterogeneous tumor cell populations. Incorporation of high-throughput single cell and spatial transcriptomic technologies to determine the effects of single or multiple cytokine and antigen combinations on specific immune subsets and T cell clones will provide a deeper understanding of the inflammatory and immunogenic signaling molecules necessary to produce different types of immune responses to help further optimize these therapeutic approaches in PDAC and potentially across other cancer types^73,74^.

Recently, neo-antigen RNA vaccines tailored to a patient’s unique mutational profile have shown promising results in early phase trials in PDAC patients in combination with anti-PD-L1 ICB and chemotherapy regimens post-surgery^50^, even resulting in persistent antigen-specific T cells peripherally in responders years later^51^. Our results suggest that multiplexed cytokine mRNA cocktails may serve as an adjuvant for antigen vaccines to further improve their efficacy by enhancing antigen presentation, innate and adaptive immune activation, and T cell memory that are typically lacking in the majority of PDAC patients but yet required for durable anti-tumor T cell responses. Whereas mRNA-lipoplexes used in these trials home preferentially to the spleen of patients that are often removed during surgery, or amphiphile-modified KRAS vaccines used in other trials home to the lymph nodes^75^, both our intratumoral as well as NP approaches direct both cytokine and antigen signals to the PDAC TME to mediate local immune responses there. Moreover, while these vaccine trials treated patients with minimal residual disease post-surgery and in combination with other chemotherapy and immunotherapy regimens, we show that combined cytokine and antigen mRNA therapy can be used to treat established and treatment naïve tumors in preclinical PDAC models, leading to tumor ablation and complete responses in 50% of mice. As work from other groups has recently shown that TAAs may outnumber mutated neo-antigens and can provide durable antigen-specific immunity against PDAC and other tumor types^73,76^, our platform using TAAs could offer an off-the-shelf approach to broaden the application of cancer vaccine strategies to more patients. Though not explored here, given that NPs deposit in the liver following systemic delivery, it would be of interest to explore whether multiplexed cytokine/antigen mRNA therapy approaches alone or in combination with other immunotherapies could be used to target late-stage metastatic disease, which >50% of PDAC patients present with at initial diagnosis and typically in the liver. Collectively, our results put forward an all-in-one cytokine and antigen mRNA immunotherapy capable of producing complete tumor responses in preclinical PDAC models with clear translational relevance.

## METHODS

### Ethical regulations

The research performed in this study complies with all ethical regulations. All mouse experiments were approved by the University of Massachusetts Chan Medical School Internal Animal Care and Use Committee (IACUC) (PROTO202000077).

### mRNA synthesis and purification

Mouse cDNA sequences encoding IL-12p70 (p40p35) (Addgene; Cat. No. 108665; gift from Nevil Singh), IL-15Rα (OriGene; Cat. No. MR226273), IL-18 (OriGene; Cat. No. MR226935), IFN-β1 (OriGene; Cat. No. MR226101), CCL5 (OriGene; Cat. No. MG227153), CXCL10 (OriGene; Cat. No. MR200291), MUC-1 (OriGene; Cat. No. MR209607), MSLN (OriGene; Cat. No. MR209539), and FOLH1 (PSMA; OriGene; Cat. No. MR216035) were cloned into a pUC19 vector backbone (a gift from Dr. L. Li at UMass Chan) that includes a T7 promoter, 5’ UTR, 3’ UTR, and 110 bp poly A tail using Gibson Assembly (New England Biolabs; Cat. No. E5510S) according to the manufacturer’s protocol.

cDNA containing pUC19 plasmids were then linearized downstream of the poly(A) signal using EcoRI restriction digestion (New England Biolabs; Cat. No R3101L). For synthesis and delivery of free (naked) mRNAs, *in vitro* transcription (IVT) was performed using T7 RNA polymerase (New England Biolabs; Cat. No E2050L) in the presence of a modified nucleotide mix, wherein uridine 5′-triphosphate (UTP) was substituted with N1-methylpseudouridine-5′-triphosphate (m¹ΨTP) (TriLink; Cat. No N-1081-1) to enhance RNA stability and reduce immunogenicity. The IVT reaction also included ATP, CTP, and GTP at equimolar concentrations. Following transcription, the RNA was capped using a 7-methylguanosine (m⁷G) cap analog (CellScript; Cat. NoC-SCCS1710) to ensure efficient translation. For synthesis of mRNAs to be encapsulated into lipid nanoparticles (NPs), IVT and post-transcriptional capping were performed using the mScript™ mRNA Production System (CellScript; Cat. No IMMY240625) according to the manufacturer’s instructions. Scramble mRNA was synthesized from the pUC19-BNTns-EGFP vector lacking a start codon to prevent translation. A poly(A) tail (110 bps) was encoded in the DNA template to produce a polyadenylated transcript. The resulting capped and polyadenylated mRNA was purified using cellulose column chromatography to remove double-stranded RNA contaminants (Messenger Bio; Cat. No EMBARK100). Synthesized and purified IVT mRNAs were analyzed by electrophoresis on a 2% agarose gel stained with ethidium bromide to verify integrity and size. A single-stranded RNA ladder (ThermoFisher Scientific, Cat. No. SM1823) was included as a molecular weight reference.

### mRNA lipid nanoparticle (NP) generation and characterization

mRNA NPs were synthesized via microfluidic co-flow focusing using a commercially available Fluigent Raydrop. MC3 (48 mol% D-Lin-MC3-DMA, Cayman Chemical, 25325) and DSPC (10 mol% 1,2-distearoyl-sn-glycero-3-phosphocholine, Avanti, 850365C) were prepared in chloroform, along with 37 mol% cholesterol and 5 mol% DMG-PEG (1,2-dimyristoyl-rac-glycero-3-methoxypolyethylene glycol-2000, Avanti, 880151P), and formulations were dried overnight to form lipid films. Films were rehydrated in 200 proof ethanol. For microfluidic synthesis, mRNA stocks were diluted in sodium acetate buffer (pH=3) at 20-200ug/mL and used as the outer aqueous phase and lipids rehydrated in ethanol were used as the inner organic phase in a microfluidic synthesis. The flow rates of the inner and outer phase were set to 12uL/min and 72uL/min, respectively. Following synthesis, mRNA NPs were dialyzed for 2h against nuclease-free PBS. Dynamic light scattering (DLS) and zeta potential were used to measure NP hydrodynamic size and surface charge, respectively, using a Malvern Zetasizer. A commercially available mRNA detection kit (Quant-it RiboGreen, Invitrogen, R11490) was used to analyze loading capacity and encapsulation efficiency following dialysis.

### Cell lines and compounds

Murine PDAC *KPC1* and *KPCY 2838c3* cell lines were generated as previously described from *KPC* and *KPCY* GEMM animals^12,49^. The *2838c3 KPCY* cell line was acquired from Dr. B. Stanger through an MTA with the University of Pennsylvania. Cancer-associated fibroblast (CAF) cell lines were derived from *KPCY* PDAC lesions and provided by Dr. J. Pitarresi. All cell lines were cultured in Dulbecco’s Modified Eagle Medium (DMEM) supplemented with 10% fetal bovine serum (FBS) and 100 IU/mL penicillin-streptomycin (P/S), and maintained at 37 °C in a humidified incubator with 5% CO₂. *KPC1* cells were cultured on tissue culture dishes pre-coated with 100 µg/mL collagen (PureCol; Advanced BioMatrix, Cat. No. 5005). All cell lines tested negative for Mycoplasma contamination.

Trametinib (S2673) and palbociclib (S1116) were purchased from Selleck chemicals for *in vitro* studies. Drugs were dissolved in DMSO (vehicle) to yield 10mM stock solutions and stored at −80^°^C, and growth media with or without drugs was changed every 2-3 days.

### mRNA transfection *in vitro*

Murine *KPC* PDAC cells, CAFs, and splenocytes were transfected with either scramble mRNA (2.5 - 3 µg) or cytokine-encoding mRNAs (0.5 µg each) using Lipofectamine MessengerMax reagent (ThermoFisher; Cat. No. LMRNA001), according to the manufacturer’s instructions.

### Animal models

All mouse experiments were approved by the University of Massachusetts Chan Medical School Internal Animal Care and Use Committee (IACUC). Mice were maintained under specific pathogen-free conditions, and food and water were provided ad libitum. Housing conditions included a 12:12 light/dark cycle, with the lights coming on at 0700 and going off at 1900 daily, a temperature range of 20-26°C, and a humidity range of 30-70%. C57BL/6 female mice were purchased from Charles River Laboratories and *P48-Cre* mice purchased from Jackson Laboratory. *Trp53^fl/fl^*and *Kras^LSL-G12D/wt^* breeding pairs were generously provided by Dr. W. Xue. For tumor transplantation studies and *ex vivo* immune assays, only female C57BL/6 mice were used, as this greatly reduced costs and complications of housing adult male animals in the same cage. For studies using *KPC* GEMM mice, both male and female mice were used. Animal sex was not considered in the study design. Tumors did not exceed the maximum tumor size of 1,500 mm^3^ permitted by the University of Massachusetts Chan Medical School IACUC.

### Pancreatic orthotopic transplant models

Orthotopic pancreatic tumors were established by transplanting *KPC1* or *KPCY 2838c3* cells into the pancreas of 8-to 12-week-old female C57BL/6 mice. *KPC1* (4 × 10⁴) or *KPCY 2838c3* (1 × 10⁵) cells were resuspended in 25 μl of Matrigel (BD) and diluted 1:1 with cold advanced DMEM/F12 medium. Mice were anesthetized using 2–3% isoflurane, and a small incision was made on the left flank to expose the pancreas. The cell suspension was injected into the tail region of the pancreas using a Hamilton syringe, and successful delivery was confirmed by the formation of a fluid bubble without leakage into the abdominal cavity. The abdominal wall was closed with absorbable Vicryl sutures (Ethicon), and the skin was secured with wound clips (CellPoint Scientific Inc.). Tumor induction and growth were monitored by ultrasound imaging. Once tumors formed (ten to sixteen days after implantation), mice were randomized into treatment groups based on tumor volume. Upon euthanasia, pancreatic tumor tissue was separated for either fixation in 10% formalin for H&E and IHC staining, or digested into a single-cell suspension for flow cytometry analysis.

### *KPC* GEMMs

*KPC* genetically engineered mouse models (GEMMs) were generated by interbreeding *Trp53^fl/fl^, Kras^LSL-G12D/wt^*, and *P48-Cre* strains on a C57BL/6 background. Male and female *KPC* mice aged 3 to 8 months were used for all experiments. Tumor progression was monitored by ultrasound imaging, and mice were enrolled and randomized into treatment groups once tumors reached approximately 30 mm³ in volume. Pancreas tumor tissue collected upon sacrifice was fixed in 10% formalin.

### Ultrasound imaging

High-resolution ultrasound imaging was conducted using the Vevo 3100 system equipped with an MS250 13–24 MHz transducer (VisualSonics). Imaging was performed to stage pancreatic tumors prior to randomization into treatment groups and to monitor tumor progression over time. Tumor volumes were quantified using Vevo LAB analysis software.

### Intratumoral and systemic mRNA dosing in mice

For intratumoral mRNA delivery, single or combinatorial cocktails of free or NP encapsulated scramble, cytokine, and antigen mRNAs were injected directly into orthotopically transplanted *KPC* tumors in mice (as verified by ultrasound imaging) 10–16 days post-transplantation. Mice were lightly anesthetized with isoflurane to minimize movement during injection. One investigator gently palpated the abdomen to locate and stabilize the tumor near the surface of the skin, while a second investigator administered the mRNA formulation directly into the tumor through the skin using a Hamilton syringe. 50µl of mRNAs diluted in Ringer’s lactate (ThermoFisher; Cat. No. J67572.AP) at varying concentrations were delivered per injection. Care was taken to avoid leakage and ensure accurate delivery into the tumor mass. Mice received either a single injection or repeated twice weekly intratumoral injections. For systemic delivery of mRNA encapsulated NPs into *KPC* GEMMs with confirmed PDAC tumors, mice received 200µl intravenous tail vein injections of empty or mRNA loaded NPs containing ∼10 µg of each mRNA weekly for two weeks or until survival endpoint. Ultrasound imaging was used to track tumor responses, and animal body weight changes and measurements of cytokines in the blood and liver were used to monitor toxicity.

### NK and CD8 depletion *in vivo*

For NK cell depletion, mice were administered intraperitoneal (i.p.) injections of an anti-NK1.1 antibody (250 μg; PK136, BioXcell; catalog no: BE0036) twice weekly. For CD8⁺ T cell depletion, mice received i.p. injections of anti-CD8 antibody (200 μg; 2.43, BioXcell; catalog no: BE0061) twice weekly. Effective depletion of NK and CD8⁺ T cells was confirmed by immunohistochemical analysis of pancreatic tumor tissue.

### Cytokine array

For *in vitro* cytokine array analysis, murine *KPC1* PDAC cells or CAFs were plated in six-well plates and either transfected with scramble or cytokine mRNA cocktails, or treated with trametinib (25nM) or palbociclib (500nM) (T/P). 24 hours post-transfection of mRNAs or 8 days post-treatment with T/P, conditioned media was collected, and cells were trypsinized and counted using a Countess II cell counter (Invitrogen). Media samples were then normalized to cell number by dilution with fresh culture media. A 75 μl aliquot of each sample was analyzed using the Mouse Cytokine/Chemokine 44-Plex Discovery Assay Array (MD44) or Mouse IL-18 Single Plex Discovery Assay Array (MDIL18) from Eve Technologies.

For cytokine array analysis of *in vivo* samples, tumor tissue, liver tissue, and blood were collected at different timepoints from *KPC* orthotopic transplant or GEMM animals following a single dose of free or NP formulated mRNAs delivered by intratumoral or intravenous injection. Tumor and liver tissues were mechanically dissociated using the MACS Dissociator (Miltenyi Biotec) prior to protein extraction. Total protein concentrations were quantified using the BCA kit (Bio-Rad; Cat. No 5000002). A total of 100 μl of each normalized sample was analyzed using the Mouse Cytokine/Chemokine 44-Plex Discovery Assay Array (MD44) or Mouse IL-18 Single Plex Discovery Assay Array (MDIL18) from Eve Technologies.

### NK and T cell migration assays

Primary NK cells and CD8⁺ T cells were isolated from the spleens of 8-to 12-week-old female C57BL/6 mice using the NK Cell Isolation Kit II (Miltenyi Biotec; Cat. No 130-115-818) and CD8a⁺ T Cell Isolation Kit (Miltenyi Biotec; Cat. No 130-104-075), respectively, according to the manufacturer’s instructions. A total of 50,000 NK or CD8⁺ T cells were diluted in serum-free DMEM supplemented with 100 IU/ml penicillin/streptomycin and seeded into the upper chamber of transwell inserts (Corning) placed in 24-well plates. Serum-free conditioned media collected 24 hours post-transfection of *KPC1* tumor cells with single or combinatorial mRNA cocktails was added to the lower chamber. After 4 hours of incubation at 37 °C in a humidified incubator with 5% CO₂, migrated cells in the lower chamber were fixed with 4% paraformaldehyde (PFA), stained with DAPI, and quantified using a Celigo imaging cytometer (Nexcelom).

### NK and T cell co-culture assays

Primary NK and CD8⁺ T cells were isolated from spleens of 8-to 12-week-old naïve female C57BL/6 mice using the NK Cell Isolation Kit II (Miltenyi Biotec; Cat. No 130-115-818) and CD8a⁺ T Cell Isolation Kit (Miltenyi Biotec; Cat. No 130-104-075), respectively, according to the manufacturer’s instructions. NK or CD8⁺ T cells were then co-cultured at a 5:1 effector-to-target ratio with *KPC1* PDAC tumor cells transfected 24 hours earlier with different single or combinatorial mRNA cocktails in RPMI media supplemented with 10% FBS, PMA (20 ng/ml, Sigma-Aldrich), Ionomycin (1 μg/ml, STEMCELL technologies), and monensin (2 μM, Biolegend) in a humidified incubator at 37°C with 5% CO_2_. Following 4 hr co-culture, IFNγ expression in NK and CD8⁺ T cells was assessed by flow cytometry analysis as described below.

### Analysis of antigen-specific CD8^+^ T cell responses *ex vivo*

To determine whether antigen mRNA vaccination could generate antigen-specific CD8^+^ T cells in PDAC tumors, mice with orthotopically transplanted *KPC1* tumors were first injected intratumorally with scramble mRNA or a 3 antigen mRNA cocktail (10 µg each of MUC-1, MSLN, and PSMA; 30 µg total). 72 hrs after vaccination, CD8^+^ T cells were purified from pancreatic tumor tissue using CD8a⁺ T Cell Isolation Kit (Miltenyi Biotec; Cat. No 130-104-075) according to the manufacturer’s instructions. In parallel, splenocytes were harvested from naïve wild-type C57BL/6 mice and pulsed *ex vivo* with either scramble mRNA (3 µg) or antigen mRNAs (1 µg each of MUC-1, MSLN, and PSMA; 3 µg total) for 24 hours. Antigen-pulsed splenocytes were then co-cultured with tumor-derived T cells at a 5:1 effector-to-target ratio for 4 hrs in RPMI media supplemented with 10% FBS, PMA (20 ng/ml, Sigma-Aldrich), Ionomycin (1 μg/ml, STEMCELL technologies), and monensin (2 μM, Biolegend) in a humidified incubator at 37°C with 5% CO_2_. Antigen-specific T cell activation was assessed by flow cytometry analysis of IFNγ and GZMB expression as described below.

### Clonogenic growth assays

Murine *KPC1* tumor cells or CAFs (5 × 10^5^ cells) were seeded in six-well plates and transfected with either scramble or cytokine-encoding mRNAs. 7 days post-transfection, adherent colonies were fixed with a solution of 1% methanol and 1% formaldehyde, stained with 0.5% crystal violet, and imaged using a digital scanner. To quantify colony formation, crystal violet–stained wells were solubilized in 100% ice-cold methanol, and absorbance was measured at 560 nm (OD₅₆₀) using a microplate reader. Background signal from blank wells was subtracted to obtain final OD values.

### Immunohistochemistry

Formalin-fixed paraffin-embedded (FFPE) tissues were prepared by overnight fixation in 10% neutral-buffered formalin, followed by standard paraffin embedding and sectioning at 5 μm thickness. Hematoxylin and eosin (H&E) staining and immunohistochemistry (IHC) were performed using established protocols. For IHC, tissue sections were deparaffinized, rehydrated through a graded ethanol series, and subjected to antigen retrieval by boiling in 10 mM citrate buffer (pH 6.0) or Tris EDTA (pH 9.0) for 15 minutes using a pressure cooker. Endogenous peroxidase activity was blocked with 3% hydrogen peroxide for 20 minutes, followed by two PBS washes.

Sections were incubated overnight at 4 °C with the following primary antibodies: CD3 (1:200; Abcam, ab5690), CD8 (1:100; eBioscience, 4SM15), NKp46 (1:100; R&D Systems, AF2225), Granzyme B (GZMB; 1:100; Abcam, ab4059), α-SMA (1:500; Abcam, ab5694), MUC-1 (1:200; Invitrogen, MA5-11202), Mesothelin (MSLN; 1:200; LS Bio, LS-C407883), and PSMA (FOLH1; 1:1000; Abcam, ab314142). The following day, sections were incubated with horseradish peroxidase (HRP)-conjugated secondary antibodies using the Vectastain Elite ABC-HRP kits specific to rat, rabbit, or goat IgG (Vector Laboratories; Cat. No.: MP-7444-15, MP-7401-50, and MP-7405-15, respectively) for 30 minutes. Signal detection was performed using 3,3′-diaminobenzidine (DAB; Vector Laboratories, SK-4105).

Stained sections were imaged using an Olympus BX41TF light microscope. Positive cells for each marker (CD3⁺, CD8⁺, NKp46⁺, GZMB⁺, SMA⁺, MUC-1⁺, MSLN⁺, and PSMA⁺) were quantified using ImageJ software by averaging counts from 5–10 randomly selected high-power fields (10× or 20× magnification) per tumor section. Tumor necrosis was assessed in H&E-stained sections by measuring the proportion of the total pancreatic tumor area composed of necrotic regions using ImageJ software. Pathological assessment of pancreatic tumors in *KPC* GEMM animals following mRNA therapy was performed in a blinded manner by GI pathologist Dr. Zhen Zhao.

### Flow cytometry analysis

To assess changes in surface MHC-I expression on *KPC1* cells cultured *in vitro*, cells were harvested 48 hours post-transfection of mRNAs by trypsinization, resuspended in PBS containing 2% FBS, and stained with anti-H-2Kb antibody (1:200; eBioscience, 12-5958-80) for 30 minutes on ice. Flow cytometry was performed on a FACSymphony A5 cytometer, and data were analyzed using FlowJo software (BD).

For *in vivo* analysis, pancreatic tumors were first isolated from mice treated with various mRNA formulations. Pancreatic tumors were then minced and enzymatically dissociated using the gentleMACS Octo Dissociator with heaters (Miltenyi Biotec) in collagenase buffer (1× HBSS with calcium and magnesium, 1 mg/ml collagenase V, and 0.1 mg/ml DNase I) using the 37C_m_TDK1_1 program. Spleens used for unstained and DAPI controls were placed in 3 ml of PBS supplemented with 2% FBS in C tubes and dissociated using program m_spleen_01 on a gentleMACS Octo dissociator with heaters (Miltenyi Biotec). Following digestion, tissues were passed through 70-μm strainers, centrifuged, and resuspended in PBS + 2% FBS. Red blood cells were lysed using ACK lysis buffer (Quality Biological).

Single-cell suspensions were stained on ice for 30 minutes with the following antibodies (dilutions in parentheses): CD45 AF700 (1:320; BioLegend, 103128), CD11b BUV395 (1:1280; BD Biosciences, 565976), CD44 BV785 (1:100; BioLegend, 103059), CD62L PE-Dazzle 594 (1:200; BioLegend, 104447), NK1.1 BV605 (1:200; BioLegend, 108739), CD3 BV650 (1:300; BioLegend, 100229), CD8 PE-Cy7 (1:400; BioLegend, 100722), CD4 PE-Cy5 (1:200; BioLegend, 100410), CD11c BV785 (1:100; BioLegend, 117335), F4/80 APC (1:200; BioLegend, 123116), MHC-II PE (1:200; BioLegend, 107608), GR-1 PE-Dazzle 594 (1:200; BioLegend, 108451), and CD103a BV711 (1:100; BD Biosciences, 748255). DAPI was used to exclude dead cells. Flow cytometry was performed using BD LSR II or FACSymphony A5 instruments, and data were analyzed using FlowJo software (BD). A representative flow cytometry gating strategy is shown in Supplementary Fig. 1.

For intracellular cytokine and Granzyme B (GZMB) staining, single cell suspensions from *in vivo* tumor samples, or NK or CD8^+^ cell *ex vivo* cultures, were incubated in RPMI-1640 supplemented with 10% FBS and penicillin/streptomycin, and stimulated with PMA (20 ng/ml), ionomycin (1 μg/ml), and monensin (2 μM) for 4 hours at 37 °C. Surface staining was performed with CD45 AF700 (1:320; BioLegend, 103128), NK1.1 BV605 (1:200; BioLegend, 108739), CD3 BV650 (1:300; BioLegend, 100229), CD8 APC-Cy7 (1:200; BioLegend, 100714), and CD4 PE-Cy5 (1:200; BioLegend, 100410), followed by fixation and permeabilization using the Foxp3/transcription factor staining buffer set (eBioscience). Intracellular cytokine staining was conducted using GZMB APC (1:100; BioLegend, 515406) and IFN-γ V450 (1:100; TONBO biosciences, 75-7311-U100) antibodies. IFN-γ and GZMB expression were analyzed in CD3⁻NK1.1⁺ NK cells and CD3⁺ CD4⁺ or CD8⁺ T cells. A representative flow cytometry gating strategy is shown in Supplementary Fig. 1.

### Statistics and Reproducibility

Statistical analyses were performed as detailed in the corresponding figure legends. Data are presented as mean ± standard error of the mean (s.e.m.), with sample sizes (n) indicating biological replicates derived from independent samples. All samples meeting the appropriate experimental criteria were included in the analyses. Some mice were excluded from flow cytometry analysis if no grossly visible tumor was present, or from ultrasound tumor volume and immunohistochemical (IHC) analyses if tumors were predominantly necrotic. Mice that died prior to flow cytometry or ultrasound timepoints were also excluded. *In vivo* experimental groups were randomized to ensure comparable tumor burdens across groups, as determined by ultrasound imaging. *In vitro* experiments were randomized at the sample allocation stage. No statistical methods were used to pre-determine sample sizes.

Statistical significance was assessed using two-sided Student’s *t*-tests, log-rank (Mantel–Cox) tests, or one-way ANOVA followed by multiple comparison corrections (Tukey’s, Dunnett’s, or Sidák’s), as appropriate. Analyses were performed using Prism 10 software (GraphPad). Data were assumed to follow a normal distribution, although this was not formally tested in all cases. A *P* value of <0.05 was considered statistically significant.

## DATA AVAILABILITY

All data supporting the findings of this study are available from the corresponding author upon request.

## CODE AVAILABILITY

No unique code was developed for this study.

## ACKNOWLEDGEMENTS

We thank B. Stanger from the University of Pennsylvania for providing *KPCY* cell lines and L. Li from UMass Chan for providing the pUC19 plasmid; G. Cottle for technical assistance; and the Ruscetti and Pitarresi laboratories for helpful suggestions and comments on the manuscript. This work was supported by funds from the Pancreatic Cancer Alliance to M.R. and J.R.P., an NIH/NCI R00 award to a M.R. (CA241110), a Pancreatic Cancer Research Program (PCARP) Idea Development Award from the Department of Defense (DoD) to J.R.P. (PA230161P1), and an NIH/NCI R00 award to J.R.P. (CA252153). C.N.P. was supported by a fellowship from the Pancreatic Cancer Alliance. N.B. was supported by a Hopper-Belmont Inspiration Award and a postdoctoral fellowship from the Pancreatic Cancer Alliance.

## AUTHOR CONTRIBUTIONS

C.N.P and M.R. conceived the study, designed research, and wrote the manuscript with assistance from all authors; K.D.D., N. B., B.M., H.K.G., H.M., L.Z., K.C.M., and L.C. designed, performed, and analyzed experiments. G.I.K., R.D., and P.U.A. synthesized NPs. Y.Q. and W.X. assisted with mRNA design and synthesis. Z.Z. provided histopathological analysis of tissue specimens. B.C. L. and J.P.R. provided intellectual input on the project. M.R. supervised the study.

---

**Extended Data Fig. 1.**
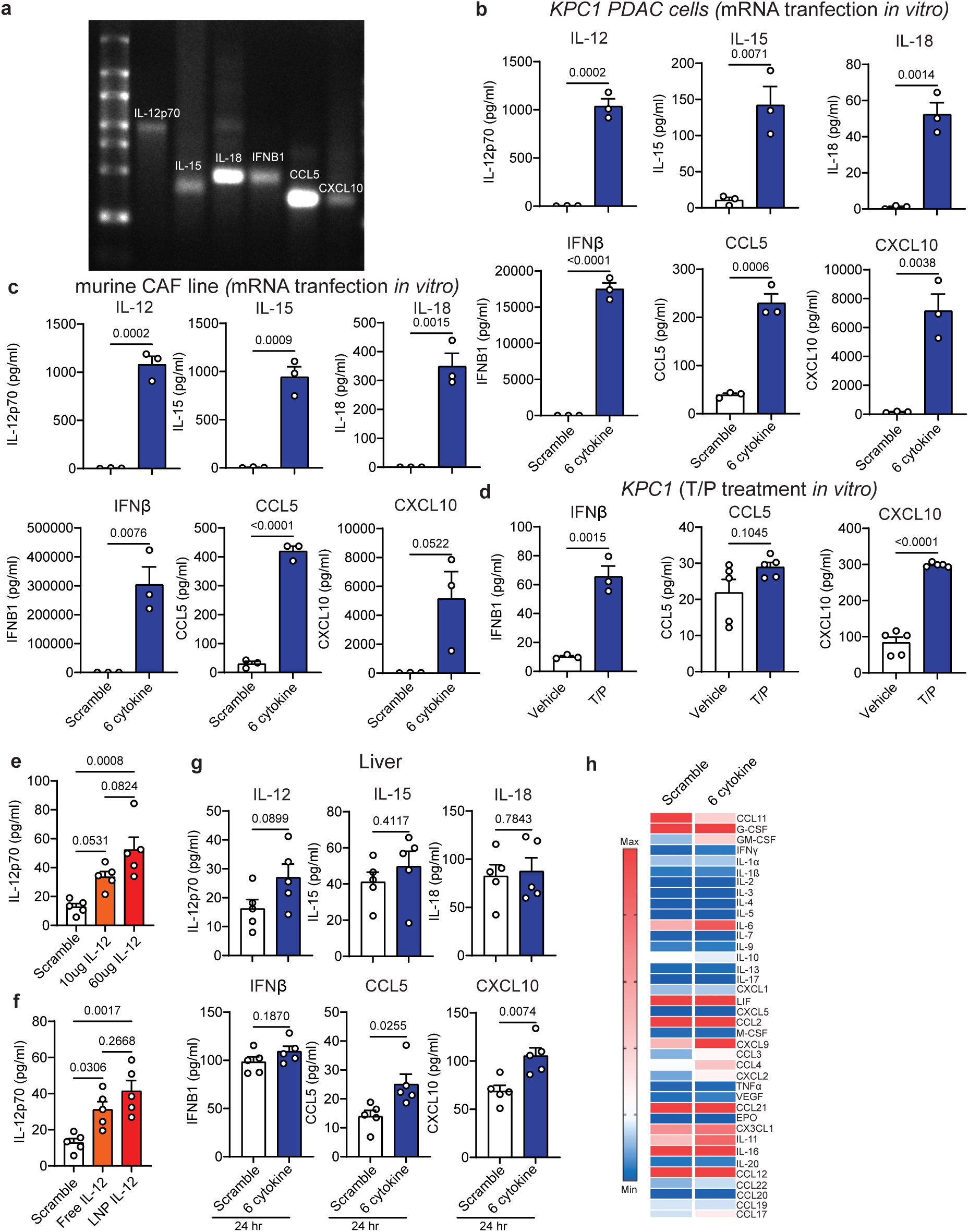
Low doses of free cytokine-encoding mRNAs can generate substantial cytokine secretion from PDAC tumor cells and fibroblasts *in vitro* and in the PDAC TME *in vivo* with limited off-target effects in the liver. **a**, Agarose gel showing cellulose-purified and capped mRNA for murine IL-12p70 (2232 bps), IL-15rα (831 bps), IL-18 (1038 bps), IFNβ1 (1088 bps), CCL5 (726 bps), and CXCL10 (756 bps) from *in vitro* transcription (IVT) as described in Fig. 1a. **b,c**, Cytokine array results of conditioned media from *KPC1* PDAC tumor cells (**b**) or cancer-associated fibroblasts (CAFs) derived from *KPCY* tumors (**c**) 24 hours after transfection with scramble mRNA (3 µg) or a 6-cytokine mRNA cocktail comprising IL-12, IL-15, IL-18, IFNβ1, CCL5, and CXCL10 (0.5 µg each; 3 µg total) *in vitro* (*n* = 3 independent samples per group). **d**, Cytokine array results of conditioned media from *KPC1* PDAC tumor cells treated with vehicle or trametinib (25nM) and palbociclib (500nM) (T/P) for 8 days (*n* = 3 independent samples per group). **e**, Cytokine array results from *KPC1* orthotopic PDAC tumors collected 24 hours post-intratumoral injection of scramble mRNA (60 µg) or different doses of IL-12 mRNA (10 or 60 µg) (*n* = 5 mice per group). **f**, Cytokine array results from *KPC1* orthotopic PDAC tumors collected 24 hours post-intratumoral injection of scramble mRNA (10 µg), free (naked) IL-12 mRNA (10 µg), or lipid nanoparticle (NP) encapsulated IL-12 mRNA (10 µg) (*n* = 5 mice per group). **g**, Cytokine array results from liver tissues collected 24 hours post-intratumoral injection of 60 µg of scramble mRNA or a 6-cytokine mRNA cocktail (10 µg each of IL-12, IL-15, IL-18, IFNβ1, CCL5, and CXCL10) (*n* = 5 mice per group) into *KPC1* PDAC tumor-bearing mice. **h**, Heatmap of cytokine array results from *KPC1* orthotopic PDAC tumors collected 96 hours post-intratumoral injection of scramble mRNA (60 µg) or a 6-cytokine mRNA cocktail (10 µg each of IL-12, IL-15, IL-18, IFNβ1, CCL5, and CXCL10; 60 µg total). Data presented as the mean of five biological replicates. *P* values in **b-d** and **g** were calculated using two-tailed, unpaired Student’s t-test, and those in **e-f** using ordinary one-way ANOVA with Tukey’s correction. Error bars are mean ± s.e.m.

**Extended Data Fig. 2.**
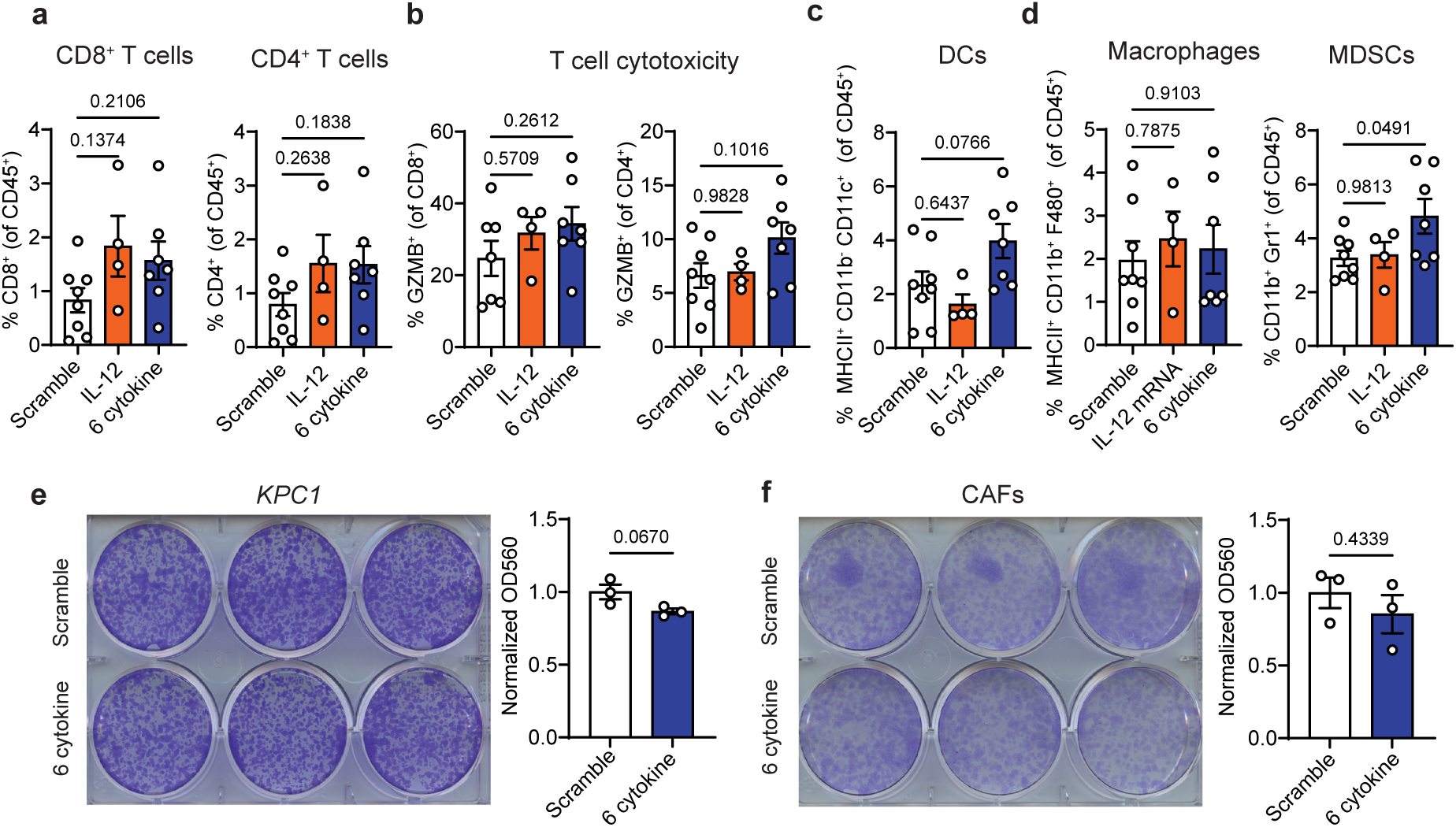
Multiplexed cytokine mRNA administration has no impact on T cell cytotoxicity or tumor cell or fibroblast viability directly in a “cold” PDAC model. **a-d**, Flow cytometry analysis of CD8^+^ and CD4^+^ T cells **(a)**, Granzyme B (GZMB)^+^ T cells **(b)**, dendritic cells (DCs) **(c)**, and MHC-II^+^ F4/80^+^ macrophages and Gr-1^+^ myeloid-derived suppressor cells (MDSCs) **(d)** in *KPC1* orthotopic PDAC tumors following repeated intratumoral injections every 3 days with scramble mRNA (60 µg), IL-12 mRNA (10 µg), or a 6-cytokine mRNA cocktail (10 µg each of IL-12, IL-15, IL-18, IFNβ1, CCL5, and CXCL10; 60 µg total) for 12 days (*n* = 4–8 mice per group). **e-f**, Representative clonogenic assay images of *KPC1* PDAC cells **(e)** and CAFs **(f)** 7 days after transfection with scramble mRNA (3 µg) or a 6-cytokine mRNA cocktail comprising IL-12, IL-15, IL-18, IFNβ1, CCL5, and CXCL10 (0.5 µg each; 3 µg total) (*n* = 3 independent samples per group). Quantification of colony formation is represented by OD560 values on the right. *P* values in **a-d** were calculated using ordinary one-way ANOVA with Dunnett’s correction, and those in **e-f** using two-tailed, unpaired Student’s t-test. Error bars are mean ± s.e.m.

**Extended Data Fig. 3.**
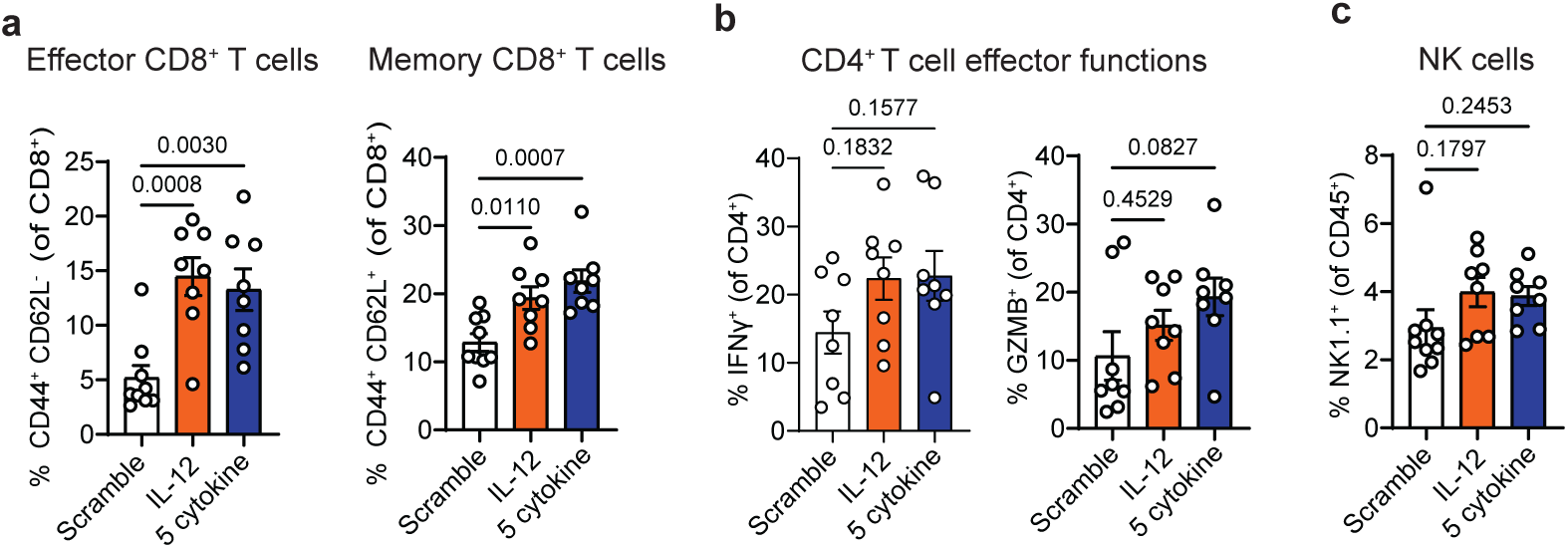
IL-12 mRNA alone or in combination with other cytokine mRNAs enhances CD8^+^ T cell effector and memory differentiation in a “hot” PDAC model expressing neo-antigens. **a-c**, Flow cytometry analysis of effector and memory CD8^+^ T cells **(a)**, CD4^+^ T cell effector functions **(b)**, and NK1.1^+^ NK cells **(c)** in *KPCY 2838c3* orthotopic PDAC tumors from mice treated with repeated intratumoral injections every 3 days of scramble mRNA (50 µg), IL-12 mRNA (10 µg), or a 5-cytokine mRNA cocktail (10 µg each of IL-12, IL-18, IFNβ1, CCL5, and CXCL10; 50 µg total) for 12 days (*n* = 8–9 independent mice per group). *P* values were calculated using ordinary one-way ANOVA with Dunnett’s correction. Error bars are mean ± s.e.m.

**Extended Data Fig. 4.**
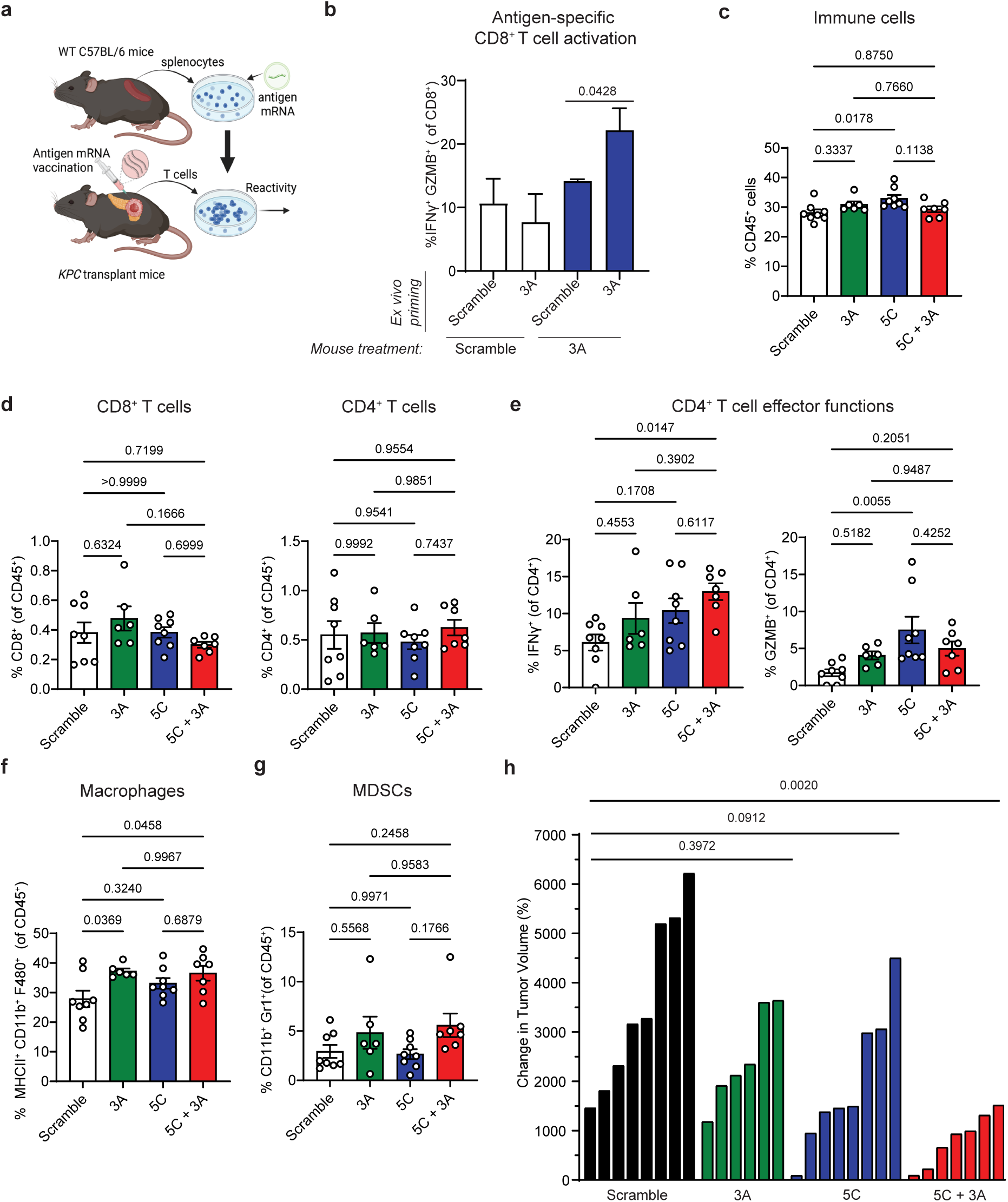
Intratumoral delivery of tumor-associated antigen mRNAs activates antigen-specific T cells and in combination with cytokine mRNAs significantly reduces tumor growth in an immunologically “cold” PDAC model. **a**, Schematic of antigen mRNA vaccination of PDAC-bearing mice and measurement of antigen-reactive T cells *ex vivo*. Splenocytes were harvested from naïve wild-type C57BL/6 mice and pulsed ex vivo with either scramble mRNA (3 µg) or antigen mRNAs (1 µg each of MUC-1, MSLN, and PSMA; 3 µg total) for 24 hours. In parallel, T cells were isolated from *KPC1* orthotopic PDAC tumors in mice that had received a single intratumoral injection of either scramble mRNA (30 µg) or 3 antigen (3A) mRNAs (10 µg each of MUC-1, MSLN, and PSMA; 30 µg total) 72 hours prior. The antigen-pulsed splenocytes were then co-cultured with tumor-derived T cells, and antigen-specific responses assessed using flow cytometry. **b**, Flow cytometry analysis of IFNγ and GZMB positivity in CD8^+^ T cells derived from *KPC1* PDAC tumors in mice vaccinated with indicated mRNAs and exposed to murine splenocytes primed with indicated mRNAs *ex vivo* as in **a** (*n* = 3-5 independent samples per group). **c-g**, Flow cytometry analysis of total CD45^+^ immune cells **(c)**, CD8^+^ and CD4^+^ T cells **(d)**, CD4^+^ T cell effector functions **(e)**, MHC-II^+^ F4/80^+^ macrophages **(f)**, and Gr-1^+^ MDSCs **(g)** in *KPC1* orthotopic PDAC tumors harvested from mice following repeated intratumoral injections every 3 days with scramble mRNA (80 µg), the 3-antigen (3A) mRNA cocktail (10 µg each of MUC-1, MSLN, and PSMA; 30 µg total), the 5-cytokine (5C) mRNA cocktail (10 µg each of IL-12, IL-18, IFNβ1, CCL5, and CXCL10; 50 µg total), or combined 5C and 3A mRNA cocktails (10 µg each of IL-12, IL-18, IFNβ1, CCL5, CXCL10, MUC-1, MSLN, and PSMA; 80 µg total) for 12 days (*n* = 6-8 mice per group). **h**, Waterfall plot of the response of *KPC1* orthotopic PDAC tumors to 12 day mRNA treatment as in **c** (*n* = 6-8 mice per group). *P* values in **b** were calculated using two-tailed, unpaired Student’s t-test, and those in **c-h** were calculated using ordinary one-way ANOVA with Tukey’s correction. Error bars are mean ± s.e.m.

**Extended Data Fig. 5.**
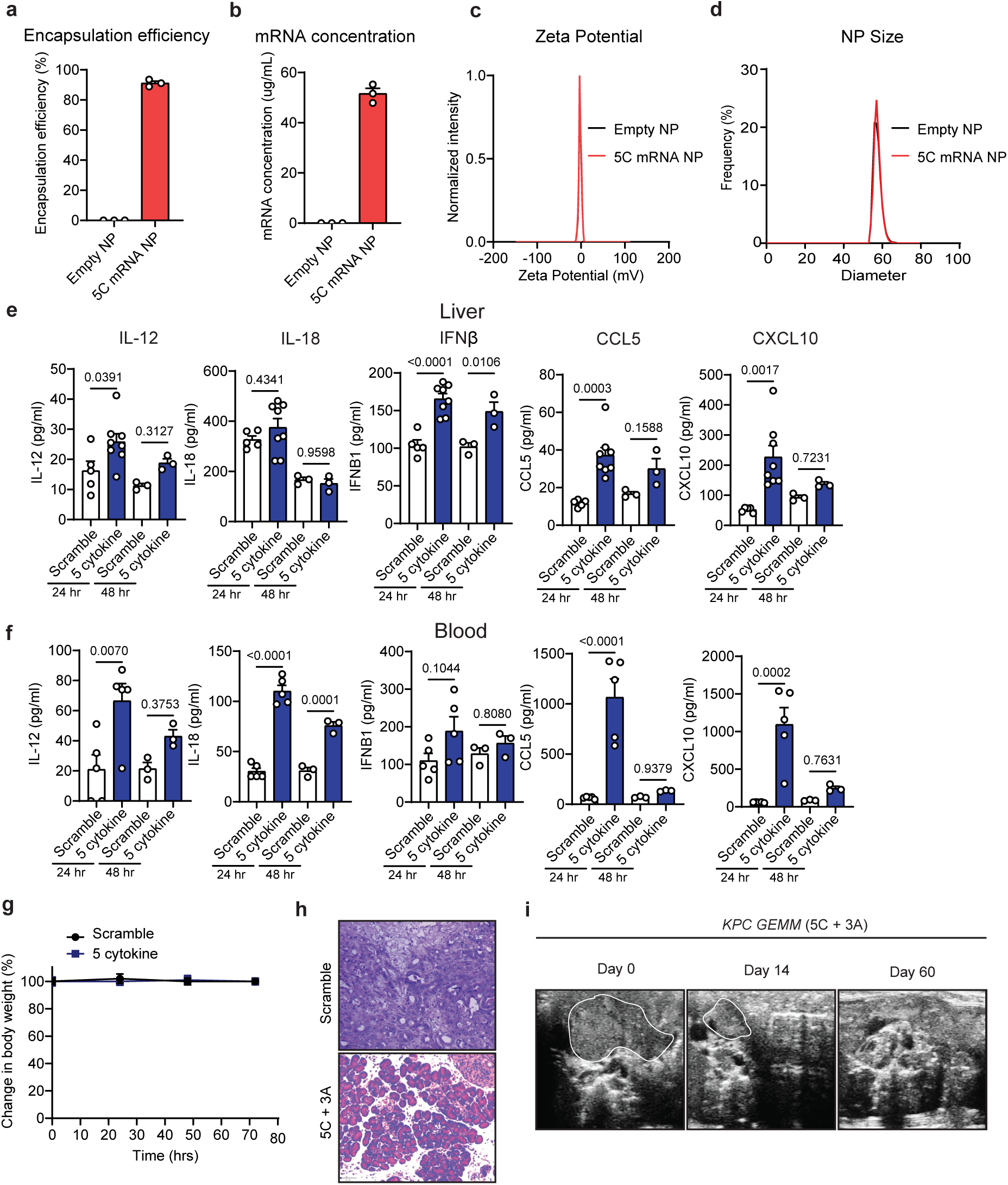
Systemic administration of NP formulated cytokine mRNAs is well-tolerated and in combination with antigen mRNAs leads to tumor regression in *KPC* GEMMs. **a**, **b**, Characterization of encapsulation efficiency **(a)** and mRNA concentration **(b)** of the 5-cytokine (5C) mRNA cocktail (IL-12, IL-18, IFNβ1, CCL5, and CXCL10) within lipid nanoparticles (NPs) (*n* = 3 independent samples per group). Empty unloaded NPs were used as controls. **c**, Measurement of NP surface charge as assessed by zeta potential. **d**, NP hydrodynamic size as assessed by dynamic light scattering (DLS). **e**,**f**, Cytokine array results from liver tissues **(e)** and blood **(f)** collected at different time points from tumor-bearing *KPC* GEMMs following tail vein injection of NPs encapsulated with scramble mRNA (50 µg) or the 5-cytokine mRNA cocktail (10 µg each of IL-12, IL-18, IFNβ1, CCL5, and CXCL10; 50 µg total) (*n* = 3–8 mice per group). **g**, Change in weight of the tumor-bearing *KPC* GEMM animals at 24-, 48-, and 72-hours following tail vein injection of mRNA NPs as in **e** (*n* = 5 mice per group). **h**, Representative H&E staining of *KPC* GEMM tumors 14 days after weekly tail vein injections of NPs encapsulated with scramble mRNA (80 µg) or the combined 5-cytokine (5C) and 3-antigen (3A) mRNA cocktail (10 µg each of IL-12, IL-18, IFNβ1, CCL5, CXCL10, MUC-1, MSLN, and PSMA; 80 µg total). Scale bar, 20 mm. **i**, Ultrasound images of a representative *KPC* GEMM tumor before treatment (Day 0) and after 14 or 60 days of weekly treatment with NPs encapsulated with the combined 5C and 3A mRNA cocktail (10 µg each of IL-12, IL-18, IFNβ1, CCL5, CXCL10, MUC-1, MSLN, and PSMA; 80 µg total). PDAC tumors are outlined in white. *P* values in **e-f** were calculated using ordinary one-way ANOVA with Sidák’s correction. Error bars are mean ± s.e.m.

## SUPPLEMENTAL FIGURE TITLES

**Supplementary Fig. 1. FACS gating strategy**

