## Supplementary figures and images for "Multiplexed cytokine and antigen mRNA administration generates durable anti-tumor immunity against pancreatic cancer"

### Supplementary Fig. 1

#### Supplementary Figure 1

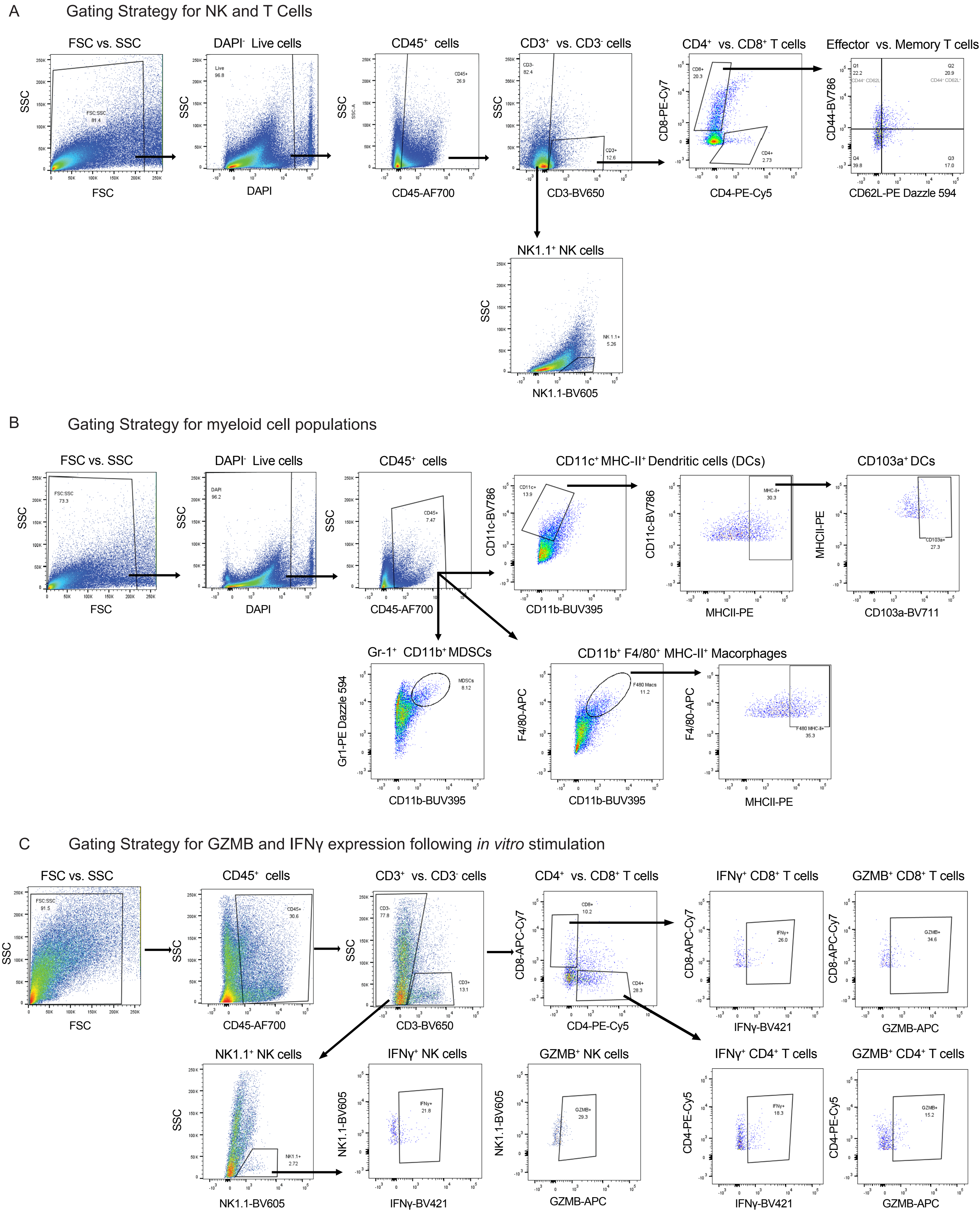
